# Acute BRCAness Induction and AR Signaling Blockage through CDK12/7/9 Degradation Enhances PARP Inhibitor Sensitivity in Prostate Cancer

**DOI:** 10.1101/2024.07.09.602803

**Authors:** Fu Gui, Baishan Jiang, Jie Jiang, Zhixiang He, Takuya Tsujino, Tomoaki Takai, Seiji Arai, Celine Pana, Jens Köllermann, Gary Andrew Bradshaw, Robyn Eisert, Marian Kalocsay, Anne Fassl, Steven P Balk, Adam S. Kibel, Li Jia

## Abstract

Current treatments for advanced prostate cancer (PCa) primarily target the androgen receptor (AR) pathway. However, the emergence of castration-resistant prostate cancer (CRPC) and resistance to AR pathway inhibitors (APSIs) remains ongoing challenges. Here, we present BSJ-5-63, a novel proteolysis-targeting chimera (PROTAC) targeting cyclin-dependent kinases (CDKs) CDK12, CDK7, and CDK9, offering a multi-pronged approach to CRPC therapy. BSJ-5-63 degrades CDK12, diminishing BRCA1 and BRCA2 expression and inducing a sustained “BRCAness” state. This sensitizes cancer cells to PARP inhibitors (PARPis) regardless of their homologous recombination repair (HRR) status. Furthermore, CDK7 and CDK9 degradation attenuates AR signaling, enhancing its therapeutic efficacy. Preclinical studies, including both *in vitro* and *in vivo* CRPC models, demonstrate that BSJ-5-63 exerts potent anti-tumor activity in both AR-positive and AR-negative setting. This study introduces BSJ-5-63 as a promising therapeutic agent that addresses both DNA repair and AR signaling mechanisms, with potential benefits for a board patient population.

## Introduction

Prostate cancer (PCa) is the second leading cause of cancer-related death in American men, trailing only behind lung cancer (*1*). Current primary treatment strategies for advanced PCa patients continue to center on androgen receptor (AR)-directed therapies, which encompass androgen deprivation therapy (ADT) and AR pathway inhibitors (ARPIs), such as enzalutamide and abiraterone. However, a significant challenge arises as patients undergoing ADT inevitably progress to a state known as metastatic castration-resistant prostate cancer (CRPC). Although ARPIs initially prove effective against CRPC, their therapeutic efficacy remains short-lived, often leading to the development of ARPI resistance (*2, 3*).

Genomic studies have revealed a variety of actionable molecular targets with underlying genomic alterations beyond the AR pathway. Notably, approximately 25% of metastatic CRPC cases exhibit genomic alterations of at least one gene involved in DNA damage response (DDR) with BRCA2 being the most frequently mutated (*4*). These alterations are associated with therapeutic vulnerabilities, particularly in the context of homologous recombination repair (HRR) deficiencies, which predict sensitivity to Poly (ADP-ribose) polymerase (PARP) inhibitors (PARPis) (*5*). PARP enzymes, especially PARP1 and PARP2, play pivotal roles in various aspects of DNA damage repair. PARPis block the repair of DNA single-strand breaks (SSBs) and result in stalled replication forks by trapping PARP1 and PARP2 on DNA breaks (*6*). This, in turn, promotes the accumulation of DNA double-strand breaks (DSBs) that HRR-deficient cells cannot repair efficiently, leading to overwhelming DNA damage and cell death. In 2020, the US FDA approved two PARPis, olaparib and rucaparib, as monotherapies for treating metastatic CRPC cases harboring BRCA1/2 mutations, capitalizing on the concept of synthetic lethality (*7–10*). The application of olaparib has expanded to include 12 additional DDR genes, the loss of which induces an HRR-deficient state, often referred to as “BRCAness” (*11*). In 2023, the FDA further approved PARPis - olaparib, niraparib, and talazoparib - for use in combination with abiraterone or enzalutamide for the treatment of metastatic CRPC cases with HRR gene mutations (*12–16*). The rationale for combining AR and PARP inhibition is that downregulation of AR-targeted DDR genes induces BRCAness (*17, 18*), while PARP1 promotes AR-mediated transcription (*19*). This combination aims to enhance therapeutic efficacy, although controversy exists regarding its use as a first-line treatment for metastatic CRPC without HHR defects (*20*). Moreover, new evidence indicates that AR does not directly regulates the transcription of DDR genes, raising doubts about whether AR inhibition alone can effectively induce a BRCAness state (*21*).

Cyclin-dependent kinase 12 (CDK12) is among the genes associated with BRCAness (*11*). Unlike other BRCAness genes directly involved in HRR, CDK12, a kinase linked to phosphorylation of the carboxy-terminal domain (CTD) of RNA polymerase II (Pol II), is essential for the transcriptional processivity of long mRNA transcripts, including those from key HRR genes, such as BRCA1 and BRCA2 (*22, 23*). CDK12 alterations, predominantly biallelic and truncating mutations, have been identified in 5-7% of PCa cases, (*24, 25*). Given that CDK12 loss-of-function mirrors the effects of BRCA1/2 mutations, CDK12 deficiency has emerged as a viable biomarker for PARPi sensitivity (*26*). Nevertheless, clinical studies have shown a lack of HRR defects and limited efficacy of PARPis in PCa patients with somatic CDK12 mutations (*24, 27–29*). In contrast, CDK12 inhibition sensitizes various cancers to PARPis by disrupting HRR processes (*30–34*), suggesting that acute CDK12 suppression through inhibitors or degraders could rapidly diminish CDK12’s modulation of HRR genes, creating a transient BRCAness state that enhances PARPi sensitivity. This approach minimizes the likelihood of cancer cells establishing feedback modulation signals to restore HRR after CDK12 genomic alterations (*35*). Consequently, a combinatorial approach involving CDK12 and PARP inhibition holds potential benefits for CRPC patients beyond their BRCA1/2 mutational status.

In addition to CDK12, CDK7 and CDK9 also have central roles in regulating Pol II by phosphorylating its CTD (*36*). Importantly, CDK7 and CDK9 are crucial for the transcriptional activation of AR. CDK7 activates MED1 by phosphorylating T1457, whereas CDK9 phosphorylates AR at Ser81, both promoting AR-mediated oncogenic transcription (*37–41*). As a master regulator of transcription, CDK7 also phosphorylates and activates CDK9 within the positive transcription elongation factor (P-TEFb) complex (*42*). The AR remains a major therapeutic target in CRPC, with MED1 serving as an intermediator between the AR and Pol II to facilitate the transcription of AR-targeted genes. Inhibition of CDK7/9 may attenuate AR signaling when direct AR targeted therapies fail, potentially overcoming enzalutamide and abiraterone resistance. Given AR’s influence on a range of DDR genes (*17, 18*), including some HRR genes, targeting CDK7/9 to inhibit AR could suppress AR-controlled HRR gene expression. This strategy may further promote the BRCAness state and increase CRPC susceptibility to PARP inhibition (*43*).

This study introduces BSJ-5-63, a novel proteolysis-targeting chimera (PROTAC) that degrades CDK12, CDK7, and CDK9, leading to the downregulation of BRCA1/2 and inhibition of AR signaling. BSJ-5-63 demonstrates potent anti-tumor effects and significantly enhances CRPC sensitivity to PARPis. The induced BRCAness state provides a therapeutic window for combining BSJ-5-63 with PARPis, offering a strategy to minimize toxicity and prevent the emergence of resistance.

## Results

### Characterization of a CDK12/7/9 degrader BSJ-5-63

We have previously developed a selective CDK12 degrader, BSJ-4-116 (fig. S1A), using the ligand of the E3 ubiquitin ligase cereblon (CRBN) (*34*). BSJ-4-116 preferentially downregulates long DDR genes such as BRCA1 and BRCA2 by enhancing intronic polyadenylation and inducing premature transcriptional termination. Consequently, BSJ-4-116 increases the sensitivity of T-cell acute lymphoblastic leukemia (T-ALL) cells to PARPis. However, BSJ-4-116 exhibited poor metabolic stability with a half-life of 2.1 minutes and intrinsic clearance of 333 µL/min/mg in hepatic microsomes, compared to the control compound, the FDA-approved multi-targeted receptor kinase inhibitor sunitinib, with a half-life of 10 minutes and intrinsic clearance of 69 µL/min/mg (table S1). Simultaneously, we have expanded our efforts by generating a series of degrader molecules. By substituting the CRBN ligand with the Von Hippel-Lindau (VHL) ligand via distinct linkers, we introduced a new CDK12 degrader, BSJ-5-63 (Fig. 1A), which showed a longer half-life of 11.1 minutes in hepatic microsomes and a lower intrinsic clearance of 62 µL/min/mg.

**Fig. 1.**
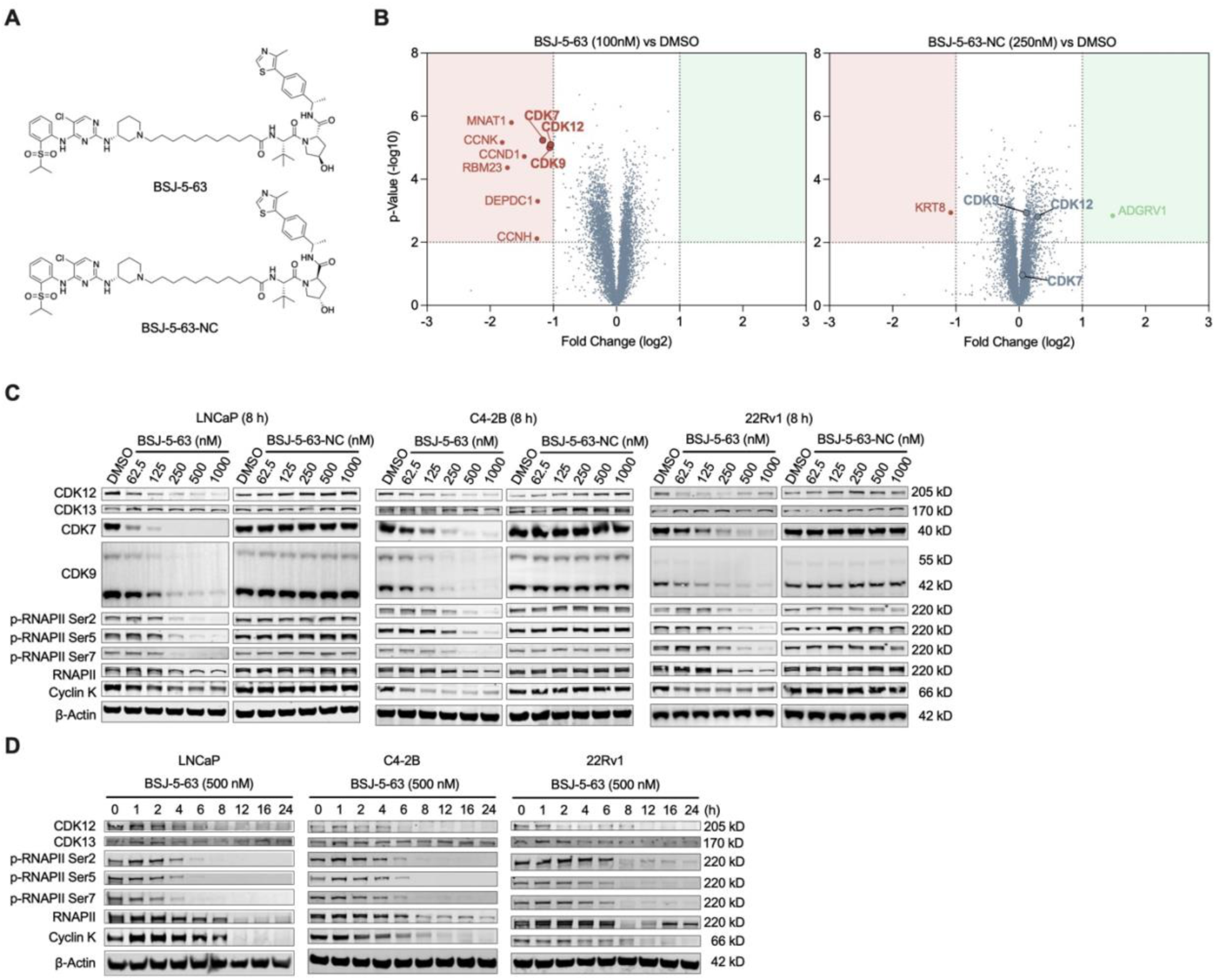
Characterization of CDK12/7/9 degrader BSJ-5-63. (A) Chemical structures of BSJ-5-63 and its negative control analog BSJ-5-63-NC. (B) Proteome-wide selectivity of BSJ-5-63. Quantitative proteomics showing the relative abundance of proteins measured by multiplexed quantitative mass spectrometry (MS)-based proteomic analysis in 22Rv1 cells treated with BSJ-5-63 (100 nM) or DMSO for 8 hours. The treatment with BSJ-5-63-NC (250 nM) was used as a control. CDK12, CDK7, and CDK9 are marked. Dotted lines indicate the threshold for statistically significant degradation of proteins. (C) Western blots of the indicated proteins in LNCaP, C4-2B and 22Rv1 cells after 8 hours of treatment with BSJ-5-63 or BSJ-5-63-NC in a dose-dependent manner. Representative blots from three independent experiments are shown. (D) Western blots of the indicated proteins in LNCaP and 22Rv1 cells after treatment with 500 nM BSJ-5-63 in a time-course manner. Representative blots from three independent experiments are shown.

To assess the specificity of BSJ-5-63 degradation, we conducted a comprehensive proteome-wide analysis using CRPC 22Rv1 cells treated with BSJ-5-63 (100 nM) for 8 hours, in parallel with the negative control BSJ-5-63-NC (250 nM), in which the stereocenter on the hydroxyproline moiety of the VHL ligand was inversed to block binding to VHL. Proteomic analysis demonstrated that BSJ-5-63 specifically degrades CDK12 and its associated partner proteins cyclin K (CCNK) while sparing CDK13 (Fig. 1B; tables S2 and S3), suggesting a potential therapeutic strategy akin to BSJ-4-116 through the suppression of DDR genes. BSJ-5-63 also degraded CDK7 and CDK9. This finding is significant in the context of PCa as CDK7 and CDK9 play pivotal roles in AR-mediated transcription and have been considered as therapeutic targets in PCa (*36, 37, 39*). Therefore, BSJ-5-63 shows potential as a treatment for PCa by targeting multiple pathways. To compare the distinct effects of the ligands CRBN and VHL in PCa cells, we examined BSJ-4-116 mediated protein degradation in 22Rv1 cells using the same proteomic analysis. Our observations revealed a notable contrast in the protein degradation profiles. While BSJ-5-63 degraded a few other proteins, including CCND1, BSJ-4-116 exhibited an increased capacity for inducing protein degradation (Fig. 1B; table S2). This raises concerns about more off-target effects when CRBN is used as a ligand in PCa cells. In addition, we did not observe degradation of CDK7 and CCNK, indicating potential differences in the therapeutic effects.

Western blot analysis further validated that BSJ-5-63 effectively induces CDK12 and CCNK degradation in PCa cell lines 22Rv1, LNCaP, and LNCaP-derived C4-2B in a dose- and time-dependent manner, whereas CDK13 protein levels remained unchanged (Fig. 1, C and D). Degradation of CDK7 and CDK9 has also been observed. CDK7 facilitates transcription initiation through phosphorylation of the CTD of RNA polymerase II (Pol II) at serines 5 and 7 (Ser5 and 7), whereas both CDK9 and CDK12 promote phosphorylation at Ser2, which is required for transcript elongation (*44*). As expected, BSJ-5-63 significantly attenuated the phosphorylation of RNA Pol II at Ser2, Ser5, and Ser7. In contrast, the negative control, BSJ-5-63-NC, had no impact on the protein levels of CDK12/7/9 or the phosphorylation of RNA Pol II. Western blot analysis of 22Rv1 cells confirmed that BSJ-4-116 induced the degradation of CDK12 and CDK9, without affecting CDK7. This resulted in a reduction of the phosphorylation of RNA Pol II at Ser 2 and 7, while Ser 5 phosphorylation remained unaffected (fig. S1C).

### BSJ-5-63 induces a BRACness state and impairs HRR activity

To investigate the impact of BSJ-5-63 on transcription, we conducted RNA-sequencing (RNA-seq) analysis of 22Rv1 cells following BSJ-5-63 treatment for 8 hours. We identified significant alterations in the expression levels of 1026 genes with BSJ-5-63 treatment as opposed to only 207 genes with the control BSJ-5-63-NC treatment (Fig. 2A; table S4). These results indicate a substantial influence on transcription as a consequence of CDK12/7/9 degradation. Consistently, gene ontology (GO) analysis revealed regulation of transcription from the RNA Pol II promoter as the most enriched downregulated pathway (Fig. 2B). This was followed by pathways associated with DSB repair via HR and the cellular response to DNA damage. Importantly, we observed significant downregulation of crucial HRR genes, including BRCA1, BRCA2, PALB2, and FANCF (Fig. 2C). RT-qPCR analysis of 22Rv1, LNCaP, and DU145 cells revealed a progressive inhibition of BRCA1 and BRCA2 mRNA expression with increasing doses and exposure times of BSJ-5-63 (Fig. 2D). Western blot analysis of LNCaP cells validated a marked reduction in BRCA1 and BRCA2 protein levels (Fig. 2E). RAD51 protein levels were also reduced, likely due to post-transcriptional regulation. Additionally, an elevation in γ-H2AX (H2AX phosphorylated at Ser 139), indicative of DSBs, was observed following BSJ-5-63 treatment. Further analyses demonstrated a time-dependent decrease in RAD51 protein levels and an increased in γ-H2AX levels across LNCaP, C4-2B, and 22Rv1 cells (fig. S2).

**Fig. 2.**
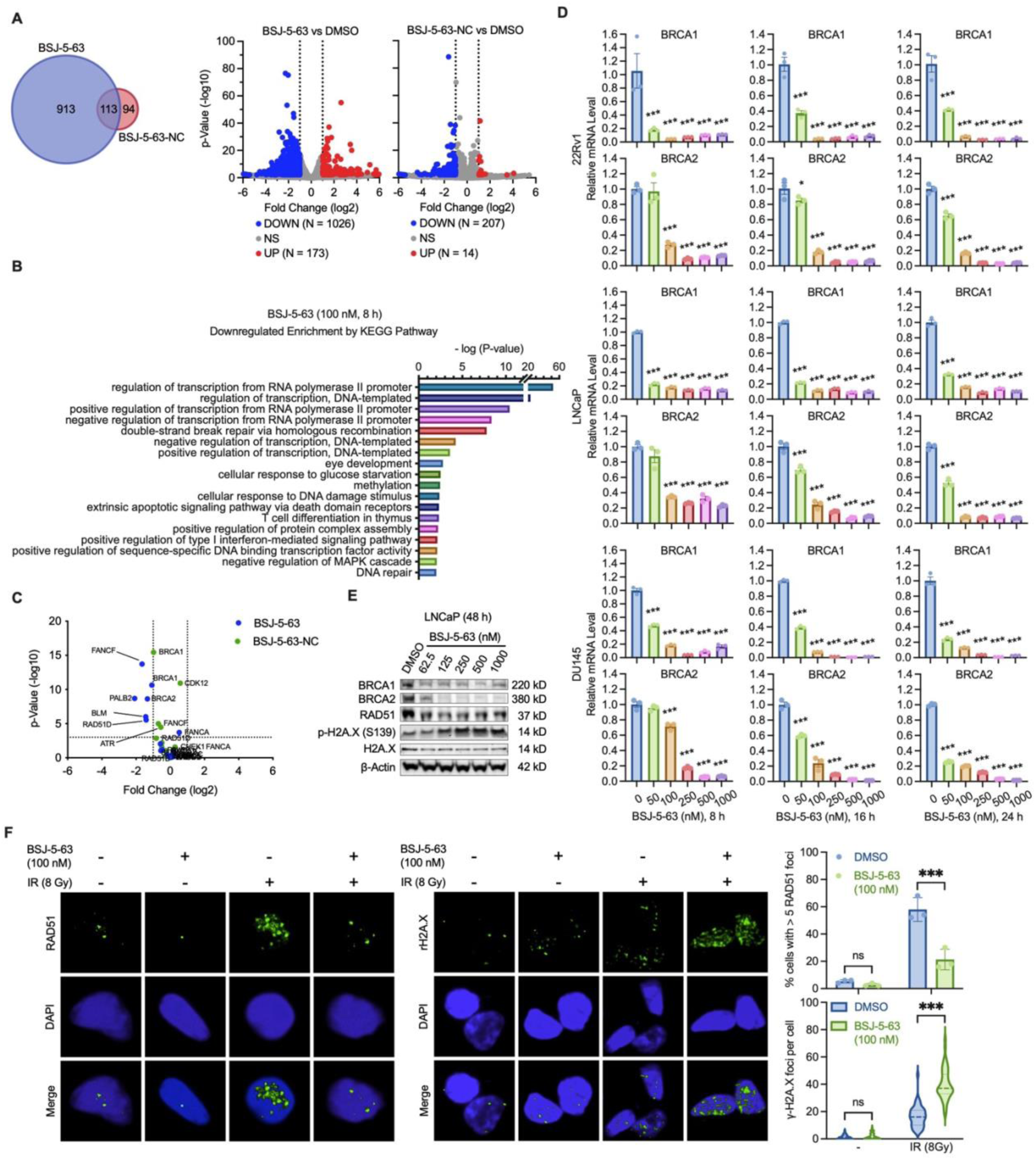
CDK12 degradation by BSJ-5-63 induces acute BRACness and impairs HRR. (A) Differentially expressed genes 8 hours after treatment with BSJ-5-63 or BSJ-5-63-NC in 22Rv1 cells. Left: Venn diagram showing the intersection of downregulated genes between BSJ-5-63 and its negative control analog BSJ-5-63-NC treatment in 22Rv1 cells. Right: Volcano plot showing differentially expressed genes after BSJ-6-63 or BSJ-5-63-NC treatment. Blue dots represent downregulated genes; Red dots represent upregulated genes. (B) Top enriched KEGG pathways in downregulated genes after BSJ-5-63 treatment in 22Rv1 cells. (C) Scatter blot showing downregulated BRACness genes after BSJ-5-63 and BSJ-5-63-NC treatment in 22Rv1 cells. (D) mRNA levels of BRCA1/BRCA2 were measured after treatment with BSJ-5-63 as indicated in 22Rv1, LNCaP, and DU145 cells by real-time RT-PCR. Data are presented as mean ± SEM (n=3). \**P* < 0.05, \*\*\**P* < 0.001; one-way ANOVA with Tukey’s multiple comparisons test. (E) Western blots of the indicated proteins in LNCaP cells after 48 hours of treatment with DMSO or BSJ-5-63 in a dose-dependent manner. Representative blots from three independent experiments are shown. (F) Representative images of immunofluorescence staining and quantification of the number of RAD51 and γ-H2AX foci in 22Rv1 cells 24 hours after BSJ-5-63 (100 nM) treatment followed by 8 Gy irradiation. Representative images from three independent experiments. \*\*\**P* < 0.001; foci were counted in at least 50 cells for each replicate under each condition (n =3). Two-way ANOVA with Tukey’s multiple comparisons test.

To determine whether BSJ-5-63 attenuates HRR function, we examined RAD51 and γ-H2AX foci formation following ionizing radiation (IR) treatment in 22Rv1 cell, with or without BSJ-5-63 (Fig. 2F). We observed an increase in RAD51 foci after IR (8 Gy), an effect that diminished in the presence of BSJ-5-63, indicating impaired HRR. Subsequently, γ-H2AX foci increased following IR, and this effect was further amplified by BSJ-5-63, suggesting an augmentation of DSBs.

These results indicate that BSJ-5-63 impairs HRR function by diminishing CDK12-mediated BRCA1/2 expression. Furthermore, treatment with BSJ-4-116 also resulted in a complete suppression of BRCA1 and BRCA2 expression (fig. S3A). This suppression was associated with a disruption of RAD51 foci formation induced by IR, indicating impaired HRR function (fig. S3B). Consequently, this disruption led to an increased occurrence of γ-H2AX foci, reflecting a higher incidence of DNA DSBs in these cells.

Next, we investigated the duration of BRCAness induced by BSJ-5-63 treatment. The 22Rv1 cells were exposed to BSJ-5-63 (100 nM) for 24 hours, followed by a medium change to eliminate the drug. We then measured BRCA1 and BRCA2 mRNA levels over time and observed that the downregulation of BRCA1 and BRCA2 expression persisted for approximately 48 and 24 hours, respectively (Fig. 3A). To explore potential synergistic effects with PARPis, we designed a sequential combination treatment for LNCaP and 22Rv1 cells, starting with a 48-hour BSJ-5-63 treatment followed by treatment with PARPis - olaparib, rucaparib, niraparib, or talazoparib (Fig. 3B). Short-term, low-dose BSJ-5-63 treatment alone had minimal effects on cell growth, suggesting lower toxicity. However, the combination therapy significantly inhibited cell growth compared to single agent treatments, as demonstrated by cell viability assays and further validated using colony formation assays in AR-positive LNCaP and 22Rv1 cells, as well as AR-negative DU145 cells (Fig. 3C). BSJ-4-116 also induced sustained BRCA1 and BRCA2 downregulation, with effects lasting for approximately 48 and 24 hours, respectively (Fig. 3D). Sequential administration of BSJ-4-116 with PARPis resulted in a significant enhancement of cell growth inhibition compared to treatment with these agents individually (Fig. 3, E and F).

**Fig. 3.**
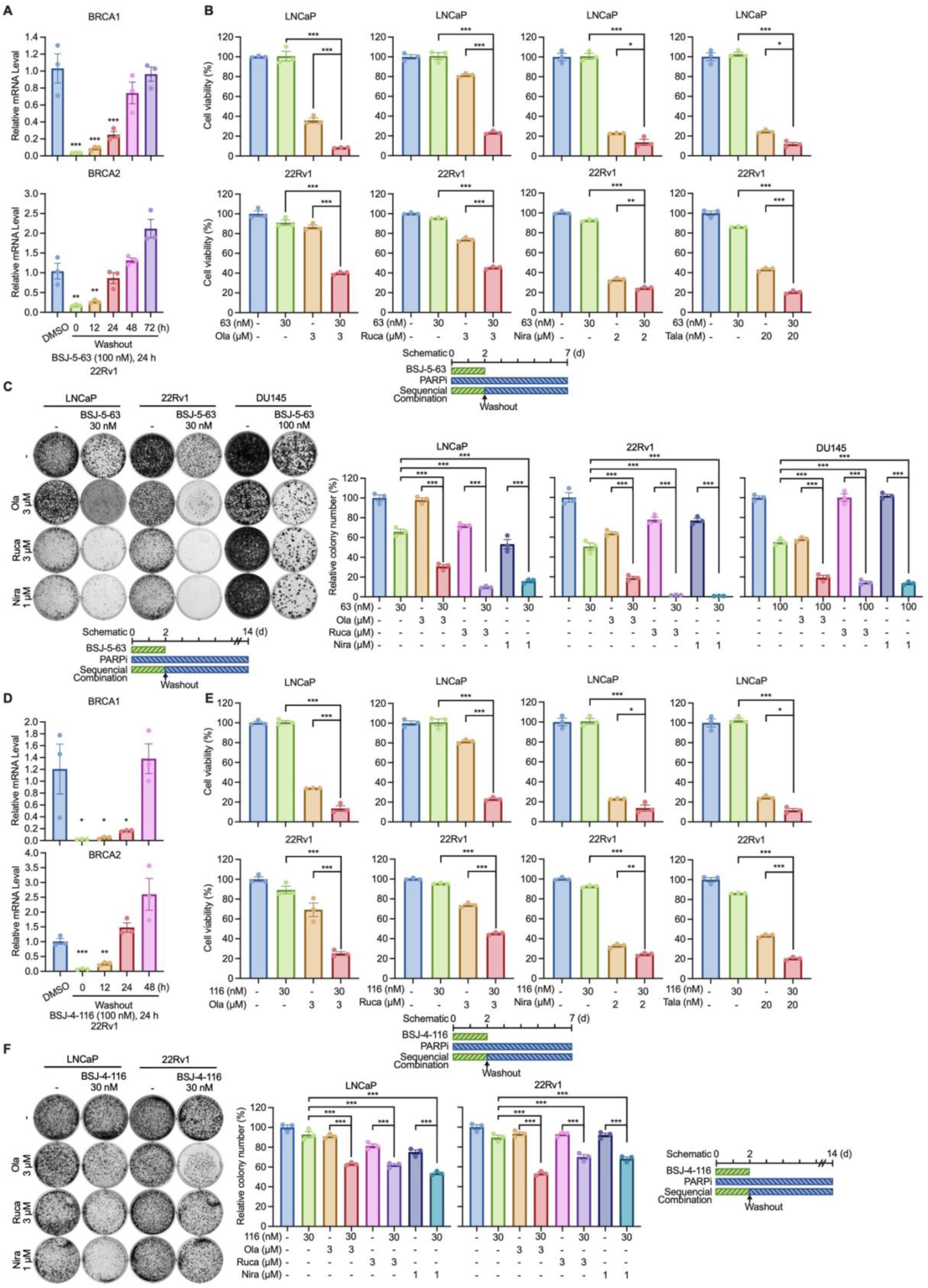
BSJ-5-63 sensitizes PCa cells to PARP inhibition through CDK12 degradation. (A) The BRCA1/2 blockage remains 24-48 hours after BSJ-5-63 washout in 22Rv1 cells. mRNA levels of BRCA1/BRCA2 were measured by real-time RT-PCR. Data are presented as mean ± SEM (n=3). \*\**P* < 0.01, \*\*\**P* < 0.001; one-way ANOVA with Tukey’s multiple comparisons test. (B) Sequential combination of BSJ-5-63 and PARPis synergistically inhibits LNCaP and 22Rv1 cell growth in a cell viability assay. BSJ-5-63 was removed from cultures through washout after 2 days of treatment, and cells were then treated by PARPis olaparib, rucaparib, niraparib or talazoparib for another 5 days as indicated. Bottom: Schematic diagram of the experimental procedure with sequential combination treatment protocol. (C) Sequential combination of BSJ-5-63 and PARPis synergistically inhibits LNCaP, 22Rv1, and DU145 colony formation. BSJ-5-63 was removed from cultures through washout after 2 days of treatment, and cells were then treated by PARPis olaparib, rucaparib, or niraparib for another 12 days as indicated. Representative colonies from three independent experiments are shown. Bottom: Schematic diagram of the experimental procedure with sequential combination treatment protocol. Data are presented as mean ± SEM (n = 3). \*\*\**P* < 0.001; one-way ANOVA with Tukey’s multiple comparisons test. (D) The BRCA1/2 blockage remains 24-48 hours after BSJ-4-116 washout. mRNA levels of BRCA1/2 were measured by real-time RT-PCR. Data are presented as mean ± SEM (n=3). \**P* < 0.05, \*\**P* < 0.01, \*\*\**P* < 0.001; one-way ANOVA with Tukey’s multiple comparisons test. (E) Sequential combination of BSJ-4-116 and PARPis synergistically inhibits LNCaP and 22Rv1 cell growth in a cell viability assay. BSJ-4-116 was removed from cultures through washout after 2 days of treatment, and cells were then treated by PARPis olaparib, rucaparib, niraparib, or talazoparib for another 5 days as indicated. Bottom: Schematic diagram of the experimental procedure with sequential combination treatment protocol. (F) Sequential combination of BSJ-4-116 and PARPis synergistically inhibits LNCaP and 22Rv1 colony formation. BSJ-4-116 was removed from cultures through washout after 2 days of treatment, and cells were then treated by PARPis olaparib, rucaparib, or niraparib for another 12 days as indicated. Representative colonies from three independent experiments are shown. Right: Schematic diagram of the experimental procedure with sequential combination treatment protocol. Data are presented as mean ± SEM (n = 3). \**P* < 0.05, \*\**P* < 0.01, \*\*\**P* < 0.001; one-way ANOVA with Tukey’s multiple comparisons test.

### Deletion of CDK12 induces a short-term BRACness and sensitize PCa cells to PARP inhibition

Many studies have demonstrated the pivotal role of CDK12 in the regulation of the expression of genes involved in HRR (*22, 45*). CDK12 deletion sensitizes cancer cells to PARP inhibition by inducing HRR defects. Interestingly, CRPC tumors with biallelic CDK12 mutations do not show a favorable response to PARP inhibition (*24*), despite the fact that most CDK12 mutations lead to truncated proteins with a loss of the kinase domain.

To further investigate this, we used CRISPR/Cas9 gene editing with two sgRNAs to delete CDK12 in 22Rv1 cells. Western blot analysis confirmed CDK12 knockout (KO), whereas CDK7 and CDK9 protein levels remained unchanged (Fig. 4A). Importantly, BRCA1 and BRCA2 protein levels were diminished in CDK12-KO cells. Further examination of RAD51 and γ-H2AX foci formation after IR treatment uncovered an increase in RAD51 foci after IR (8Gy) in CDK12-intact AAVS1 cells, a phenomenon not observed in CDK12-KO cells, indicating a HRR defect (Fig. 4B). Moreover, we observed a slightly increase in the formation of γ-H2AX foci in CDK12-intact cells after IR. This effect was dramatically amplified in CDK12-KO cells, suggesting the augmentation of DNA DSBs. CDK12-KO cells exhibited significantly enhanced sensitivity to olaparib in both the cell viability and colony formation assays (Fig. 4, C and D). Interestingly, CDK12-KO cells displayed reduced sensitivity to BSJ-5-63, implying that CDK12 is a key target of BSJ-5-63 (Fig. 4, E and F). The same results were observed when CDK12-KO cells were treated with BSJ-4-116 (fig. S4, A and B). Additionally, CDK12-KO 22Rv1 cells exhibited marked sensitivity to olaparib in xenograft studies, accompanied by a reduction in tumor weight (Fig. 4, G and H). Cell death was observed in CDK12-KO tumors using hematoxylin and eosin (H&E) staining (Fig. 4I). However, in prolonged culture under regular media conditions, the initially observed olaparib sensitivity of CDK12-KO cells was lost. We found that BRCA1 and BRCA2 expression levels were partially restored after 48 and 100 passages, indicating the potential compensation of CDK12 function through a yet to define mechanism (Fig. 4J). Taken together, our findings suggest that the acute suppression of CDK12 by genetic and pharmacological means sensitizes PCa cells to PARP inhibition. However, the long-term culture of PCa cells with CDK12 deletion suggests the possibility of compensatory mechanisms may develop over time, consistent with clinical observations in CDK12-mutant CRPC, and warrants further investigation.

**Fig. 4.**
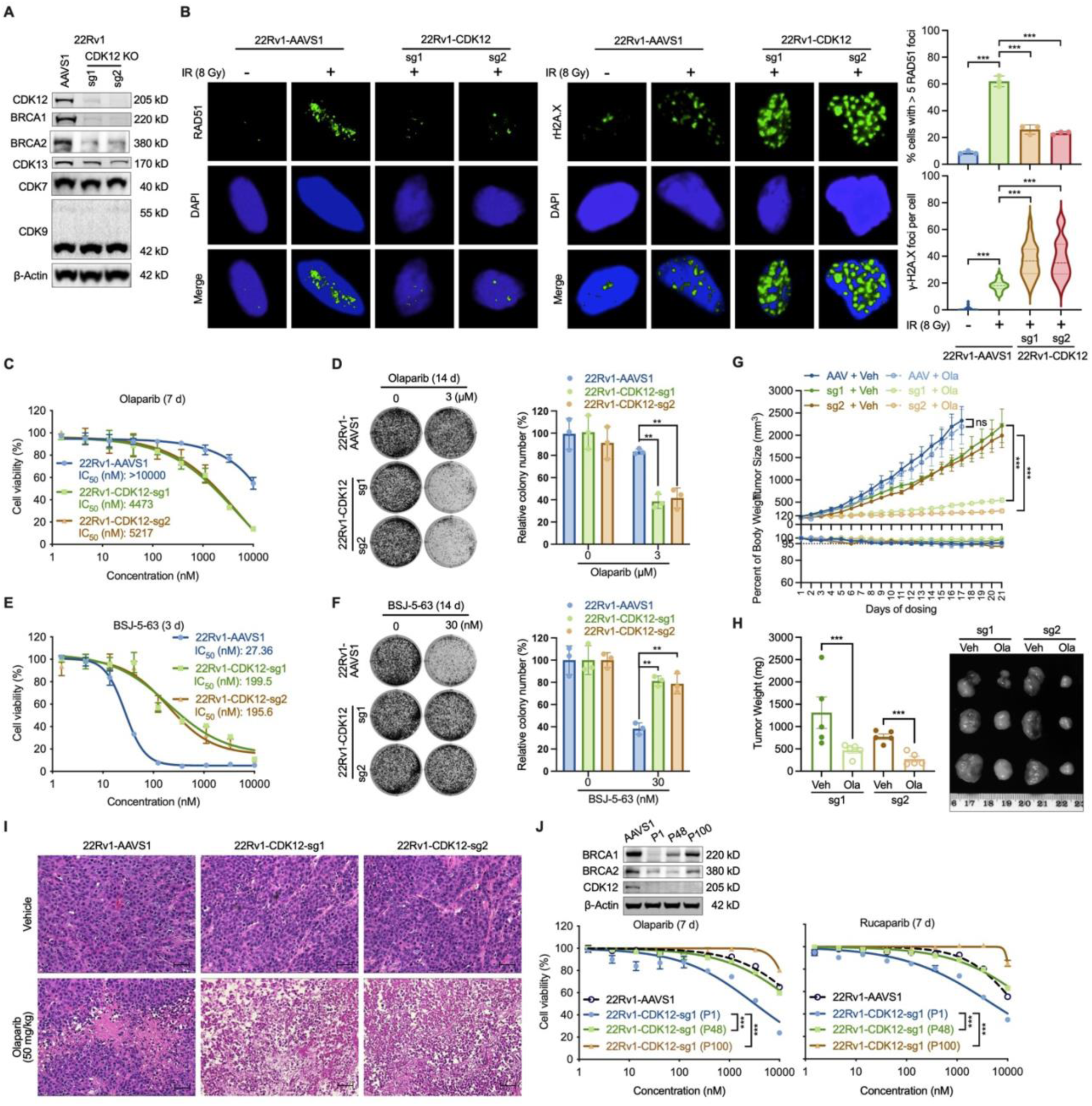
CDK12 KO impairs HRR and sensitizes PCa cells to PARP inhibition. (A) Western blots showing CDK12 KO using two different sgRNA (sg1 and sg2) in 22Rv1 cells, with the corresponding control using AAVS1 sgRNA. Representative blots from three independent experiments are shown. (B) Representative images of immunofluorescence staining and quantification of the number of RAD51 and γ-H2AX foci in CDK12-KO 22Rv1 cells after 8 Gy irradiation, with the corresponding control AAVS1-KO cells. Representative images from three independent experiments. \*\*\**P* < 0.001; foci were counted in at least 50 cells for each replicate under each condition (n =3). One-way ANOVA with Tukey’s multiple comparisons test. (C) CDK12 KO significantly sensitized 22Rv1 to olaparib in a cell viability assay. (D) CDK12 KO significantly sensitized 22Rv1 to olaparib in a colony formation assay. Representative colonies from three independent experiments are shown. Data are presented as mean ± SEM (n = 3). \*\*\**P* < 0.001; one-way ANOVA with Tukey’s multiple comparisons test. (E) CDK12 KO reduced the antiproliferation effect of BSJ-5-63 in a cell viability assay. (F) CDK12 KO reduced the antiproliferation effect of BSJ-5-63 in a colony formation assay. Representative colonies from three independent experiments are shown. Data are presented as mean ± SEM (n = 3). \*\*\**P* < 0.001; one-way ANOVA with Tukey’s multiple comparisons test. (G) CDK12 KO significantly enhanced the *in vivo* antitumor efficacy of olaparib. Top: Tumor volume curve. Bottom: Body weight curve of the mice. Data are presented as mean ± SEM (n = 5 per group). ns, not significant, \*\*\**P* < 0.001, one-way ANOVA with Tukey’s multiple comparisons test. (H) Tumor weights of CDK12-KO 22Rv1 xenografts dissected on Day 21 of drug treatment with representative photos. Data are presented as mean ± SEM (n = 5 per group). \*\*\**P* < 0.001, one-way ANOVA with Tukey’s multiple comparisons test. (I) H&E staining showing cell death after olaparib treatment in CDK12-KO 22Rv1 tumors compared with AAVS1-KO control. Scale bar: 50 μm. (J) The CDK12-KO 22Rv1 cells become resistant to PARPis after long-term passaging, compared with the first generation. Data are presented as mean ± SEM (n = 3). \*\*\**P* < 0.001; one-way ANOVA with Tukey’s multiple comparisons test. Top: Western blots showing BRCA1/2 expression levels. Representative blots from three independent experiments are shown.

### BSJ-5-63 blocks AR signaling through inhibiting MED1 and AR phosphorylation

Our proteomic analyses revealed that BSJ-5-63 effectively degrades CDK7 and CDK9, which are crucial kinases in AR-mediated transcription by phosphorylating MED1 and AR (*37–41*). Upon dihydrotestosterone (DHT) (10 nM) treatment, Western blot analysis of AR-positive LNCaP cells demonstrated increased phosphorylation of MED1 at Thr1457 and AR at Ser81, which was completely abolished by BSJ-5-63 (Fig. 5A). As a result, the levels of PSA, which is encoded by the AR target gene KLK3 and induced by DHT, were reduced. We further confirmed that CDK7 knockdown significantly reduced MED1 phosphorylation at Thr1457 in LNCaP cells (Fig. 5B), an effect that was not observed with CDK9 and CDK12 KO. Conversely, CDK9 KO reduced AR phosphorylation at Ser81 in LNCaP cells (Fig. 5C). RT-qPCR revealed a dose-dependent reduction in the mRNA levels of the AR target genes KLK3 and TMPRSS2 in LNCaP and 22Rv1 cells (Fig. 5D).

**Fig. 5.**
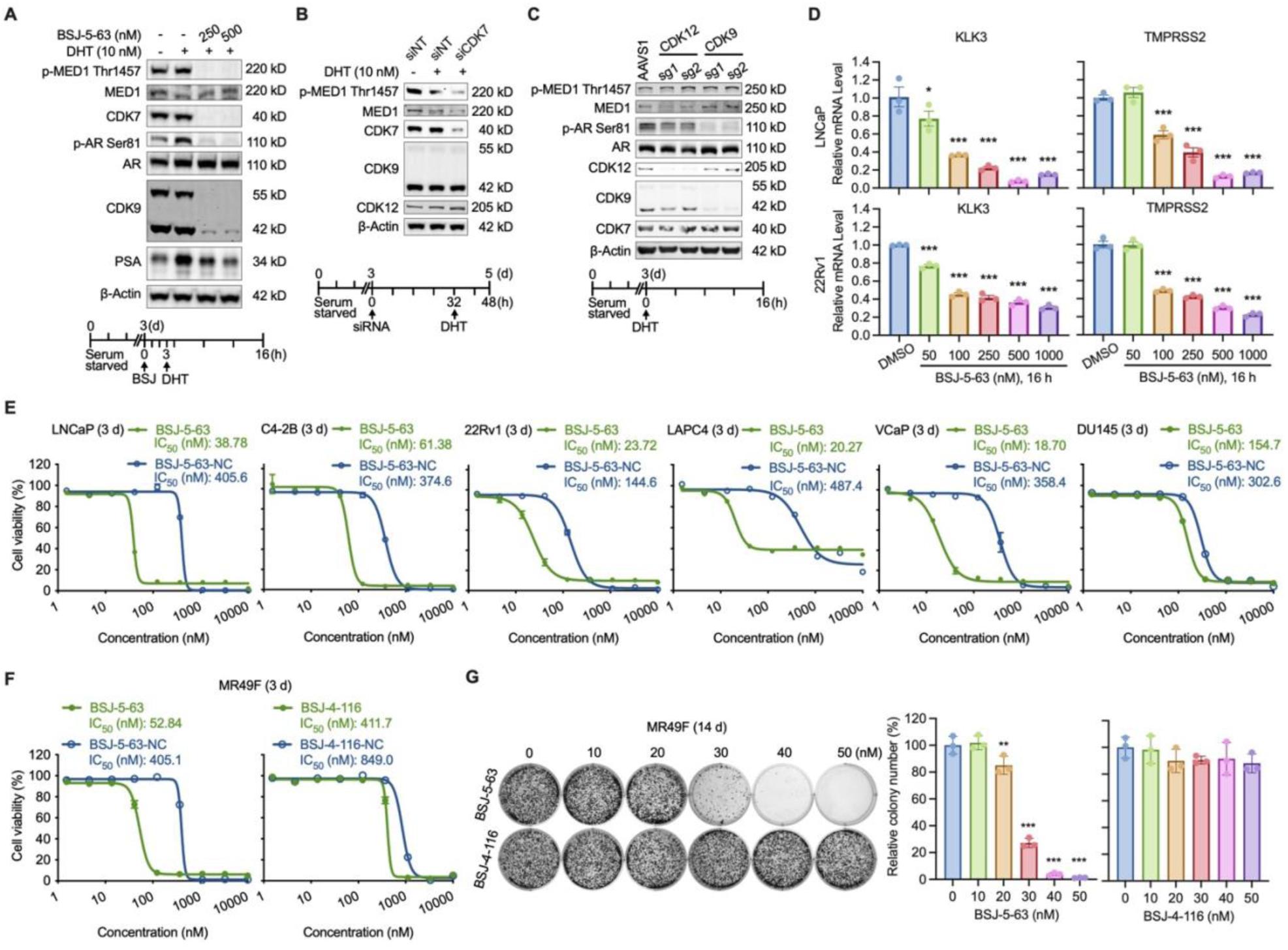
CDK7/9 degradation by BSJ-5-63 blocks MED1/AR–mediated transcription and PCa cell growth. (A) Western blot showing abolished MED1/AR phosphorylation and reduced PSA expression through CDK7/9 degradation after BSJ-5-63 treatment. LNCaP cells were grown in CSS-containing media for 3 days. Schematic of drug treatment is shown (Bottom). (B) Western blot showing reduced phosphorylation of MED1 at T1457 after CDK7 knockdown. Schematic of drug treatment is shown (Bottom). (C) Western blot showing reduced phosphorylation of AR at Ser81 after CDK9 KO. The phosphorylation of MED1 at Thr1457 remains unchanged after CDK12 or CDK9 KO. Schematic of drug treatment is shown (Bottom). (D) The expression of AR target genes significantly downregulated 16 hours after BSJ-5-63 treatment in LNCaP and 22Rv1 cells. mRNA levels of KLK3 and TMPRSS2 were measured by real-time RT-PCR. Data are presented as mean ± SEM (n=3). \*\*\**P* < 0.001; one-way ANOVA with Tukey’s multiple comparisons test. (E) Dose-response curves for LNCaP, C4-2B, 22Rv1, LAPC4, VCaP, and DU145 cells treated with BSJ-5-63 or BSJ-5-63-NC at indicated dose range for 72 hours. Data are presented as mean ± SEM (n = 3). (F) Cell viability assay showing the inhibition of MR49F cell growth after treatment with BSJ-5-63, but not BSJ-4-116. Dose-response curves for MR49F cells treated with BSJ-5-63, BSJ-5-63-NC, BSJ-4-116, or BSJ-4-1116-NC at indicated dose range for 72 hours. Data are presented as mean ± SEM (n = 3). (G) Colony formation assay showing inhibition of MR49F colony growth after treatment with BSJ-5-63, but not BSJ-4-116. Representative colonies from three independent experiments are shown. Data are presented as mean ± SEM (n = 3). \*\**P* < 0.01, \*\*\**P* < 0.001; one-way ANOVA with Tukey’s multiple comparisons test.

We evaluated the inhibitory effects of BSJ-5-63 monotherapy on PCa cell growth *in vitro*. BSJ-5-63 effectively inhibited the growth of PCa cells at nanomolar concentrations, as demonstrated by cell viability assays (Fig. 5E). The AR-positive LNCaP, C4-2B, 22Rv1, LAPC4, and VCaP cells were more sensitive to BSJ-5-63 than the AR-negative DU145 cells. This increased sensitivity was confirmed through colony formation assays in LNCaP, C4-2B, and 22Rv1 cells compared to DU145 cells (fig. S5). These results indicate that BSJ-5-63 is more effective as a monotherapy for treating AR-positive PCa cells, likely due to its ability to degrade CDK7 and CDK9. Notably, The IC_50_ values for BSJ-5-63 in five AR-positive PCa cell lines (LNCaP, C4-2B, 22Rv1, LAPC4, and VCaP) were measured as 38.78 nM, 61.38 nM, 23.72 nM, 20.27 nM, and 18.7 nM, respectively. This represents a substantial potency increase, ranging from 6.1 to 24-fold, when compared to BSJ-5-63-NC, which had IC_50_ values of 405.6 nM, 374.6 nM, 144.6 nM, 487.4 nM, and 358.4 nM, respectively. These results indicate that the growth inhibitory effect extends beyond CDK7/9 enzymatic inhibition, with protein degradation emerging as a pivotal factor in this context. Comparable outcomes were observed when PCa cells were treated with BSJ-4-116 (fig. S6). Furthermore, MR49F, an enzalutamide-resistant AR-positive PCa cell line derived from LNCaP cells (*46*), exhibited increased sensitivity to BSJ-5-63 compared to BSJ-5-63-NC and BSJ-4-116, as demonstrated by cell viability and colony formation assays (Fig. 5, F and G). The IC_50_ showed a one-log magnitude difference, supporting BSJ-5-63 as a viable therapy in cases of direct AR-targeting failure.

### BSJ-5-63 inhibits prostate tumor growth *in vivo* and synergizes with olaparib

Our *in vitro* studies revealed that BSJ-5-63 could inhibit the growth of both AR-positive and AR-negative PCa cells when used in conjunction with PARPis. This inhibition is achieved through the degradation of CDK12, leading to a BRCAness state. Alternatively, in AR-positive cells, BSJ-5-63 inhibits growth by targeting CDK7 and CDK9 for degradation, thus attenuating AR-mediated transcription. This finding prompted us to investigate the potential efficacy of this degrader in an *in vivo* setting. To begin our *in vivo* exploration, we conducted a comparative analysis of the pharmacokinetics (PK) of BSJ-5-63 in mice, in contrast to BSJ-4-116. BSJ-5-63 had a longer half-life (2.98 hours) compared to BSJ-4-116 (1.12 hours), indicating a slower clearance from the body (table S5). It also had a higher maximum plasma concentration (Cmax) and a significantly larger area under the curve (AUC), suggesting a prolonged and potentially more potent effect. These data support the *in vivo* use of BSJ-5-63.

Next, we examined the pharmacodynamics of BSJ-5-63 and its mechanism of action in mice. Western blot analysis of 22Rv1 xenograft tissues revealed the degradation of CDK12, CDK7, and CDK9 following intraperitoneal injection at a dose of 50mg/kg (Fig. 6A). Furthermore, we examined the efficacy of BSJ-5-63 as a single agent in the 22Rv1 xenograft model. Our results demonstrated that BSJ-5-63 significantly inhibited the growth of 22Rv1 tumors, whether administered daily or every three days, at a 50 mg/kg dosage (Fig. 6B). Tumor weight was significantly reduced (Fig. 6C), and there was an increased in the number of apoptotic cells after BSJ-5-63 treatment (Fig. 6D). However, it is noteworthy that daily administration resulted in approximately 10% weight loss in mice, suggesting potential toxicity associated with prolonged treatment.

**Fig. 6.**
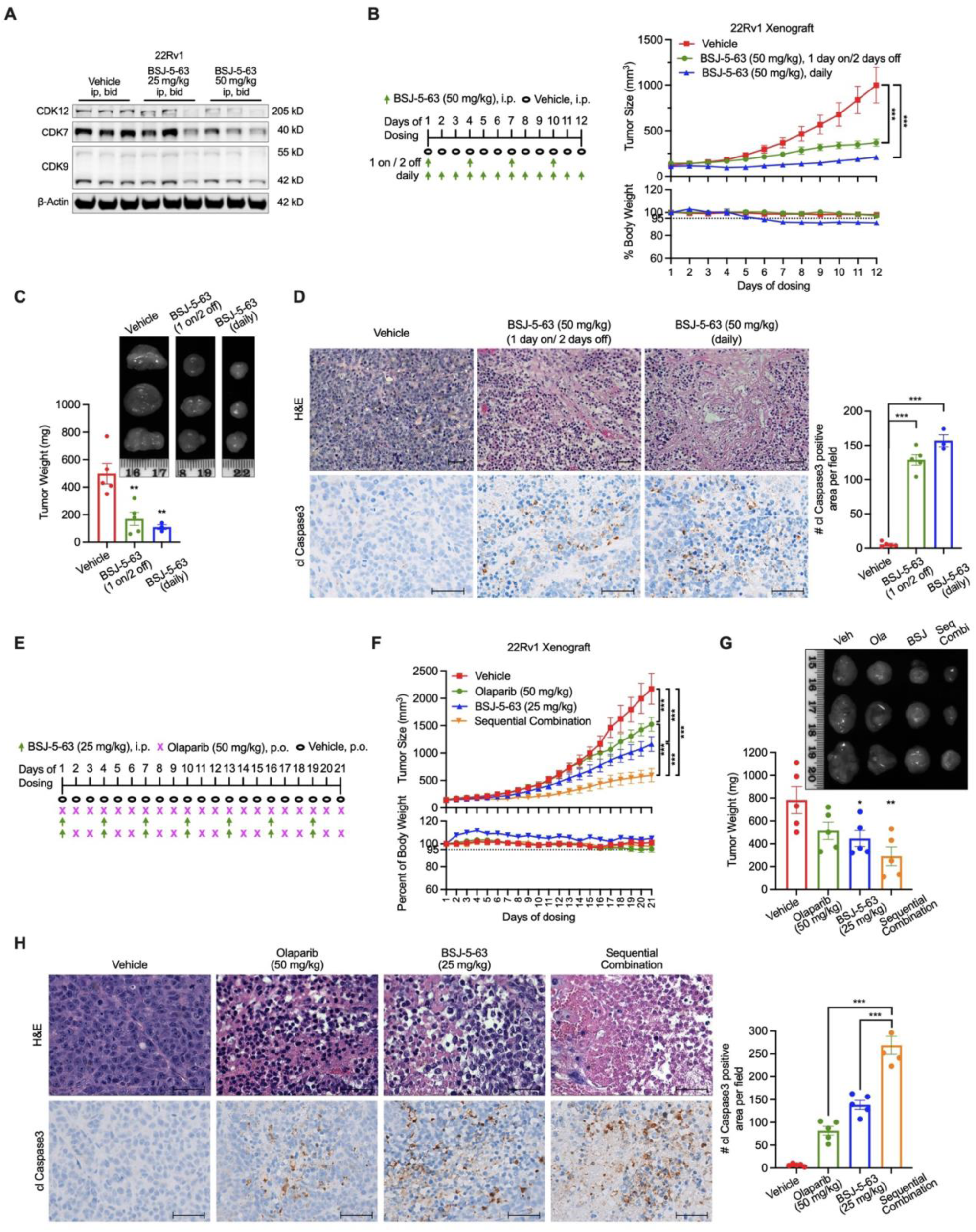
BSJ-5-63 inhibits the growth of AR-positive 22Rv1 xenografts as a single agent or in combination with olaparib. (A) Western blot analysis of the expression of CDK12/7/9 in 22Rv1 tumors from mice sacrificed 3 days after BSJ-5-63 treatment as indicated. Representative blots from three animals of each experimental group are shown. (B) The antitumor efficacy of BSJ-5-63 using as a single agent in 22Rv1 xenograft model. Left: Schematic of the BSJ-5-63 treatment protocol. Right: Tumor volume curve and Body weight curve of the mice. Data are presented as mean ± SEM (n = 3 or 5 per group). \*\*\**P* < 0.001. (C) Tumor weights of 22Rv1 xenografts dissected on Day 12 of drug treatment with representative photos. Data are presented as mean ± SEM (n = 3 or 5 per group). ***P < 0.001. (D) Immunohistochemistry analysis and H&E staining showing significantly increased apoptosis in 22Rv1 tumors after treatment with BSJ-5-63. Scale bar: 50 μm. Right: Quantification of cleaved (cl) Caspase3 staining in the xenografts. Data were presented as mean ± SEM (n = 3 or 5 per group). \*\*\**P* < 0.001 significantly different from vehicle. (E) Schematic of sequential combination treatment protocol for 22Rv1 xenograft. (F) The *in vivo* antitumor efficacy of sequential combination of BSJ-5-63 and olaparib in 22Rv1 xenograft model. Top: Tumor volume curve. Bottom: Body weight curve of the mice. Data are presented as mean ± SEM (n = 5 per group). \*\*\**P* < 0.001; one-way ANOVA with Tukey’s multiple comparisons test. (G) Tumor weights of 22Rv1 xenografts dissected on Day 21 of drug treatment with representative photos. Data are presented as mean ± SEM (n = 5 per group). \*\*\**P* < 0.001. (H) Immunohistochemistry analysis and H&E staining showing significantly increased apoptosis in 22Rv1 tumors after sequential combination of BSJ-5-63 and olapaerib. Scale bar: 50 μm. Right: Quantification of cl Caspase3 staining in the xenografts. Data were presented as mean ± SEM (n = 5 per group). \*\*\**P* < 0.001; one-way ANOVA with Tukey’s multiple comparisons test.

Subsequently, we developed a sequential therapy approach, involving a one-day treatment with BSJ-5-63 at a dosage of 25 mg/kg followed by a two-day olaparib (50 mg/kg) treatment, in comparison with treatments involving the vehicle, BSJ-5-63, and olaparib used as single agents (Fig. 6E). When administered as a monotherapy, both olaparib and BSJ-5-63 exhibited partial inhibition of 22Rv1 tumor growth (Fig. 6F). In contrast, the combination therapy had a significantly greater impact on tumor growth without causing a decrease in mouse body weight. This enhanced efficacy was further confirmed by measuring tumor size and weight post-treatment (Fig. 6G). Additionally, the number of apoptotic cells notably increased after treatment with olaparib or BSJ-5-63, and this effect was significantly intensified when the two agents were used in combination (Fig. 6H).

We then turned our attention to the AR-negative DU145 xenograft model and implemented a sequential therapy consisting of two days of BSJ-5-63 (25 mg/kg) followed by two days of olaparib (50 mg/kg) (Fig. 7A). After treatment with olaparib or BSJ-5-63 alone, we observed only partial inhibition of tumor growth (Fig. 7B). However, the combination therapy effectively halted tumor growth without any adverse effects on the body weight of the mice. Further analysis of tumor size, weight, and detection of apoptotic cells provided strong evidence supporting the synergistic effects of the combination therapy (Fig. 7, C and D).

**Fig. 7.**
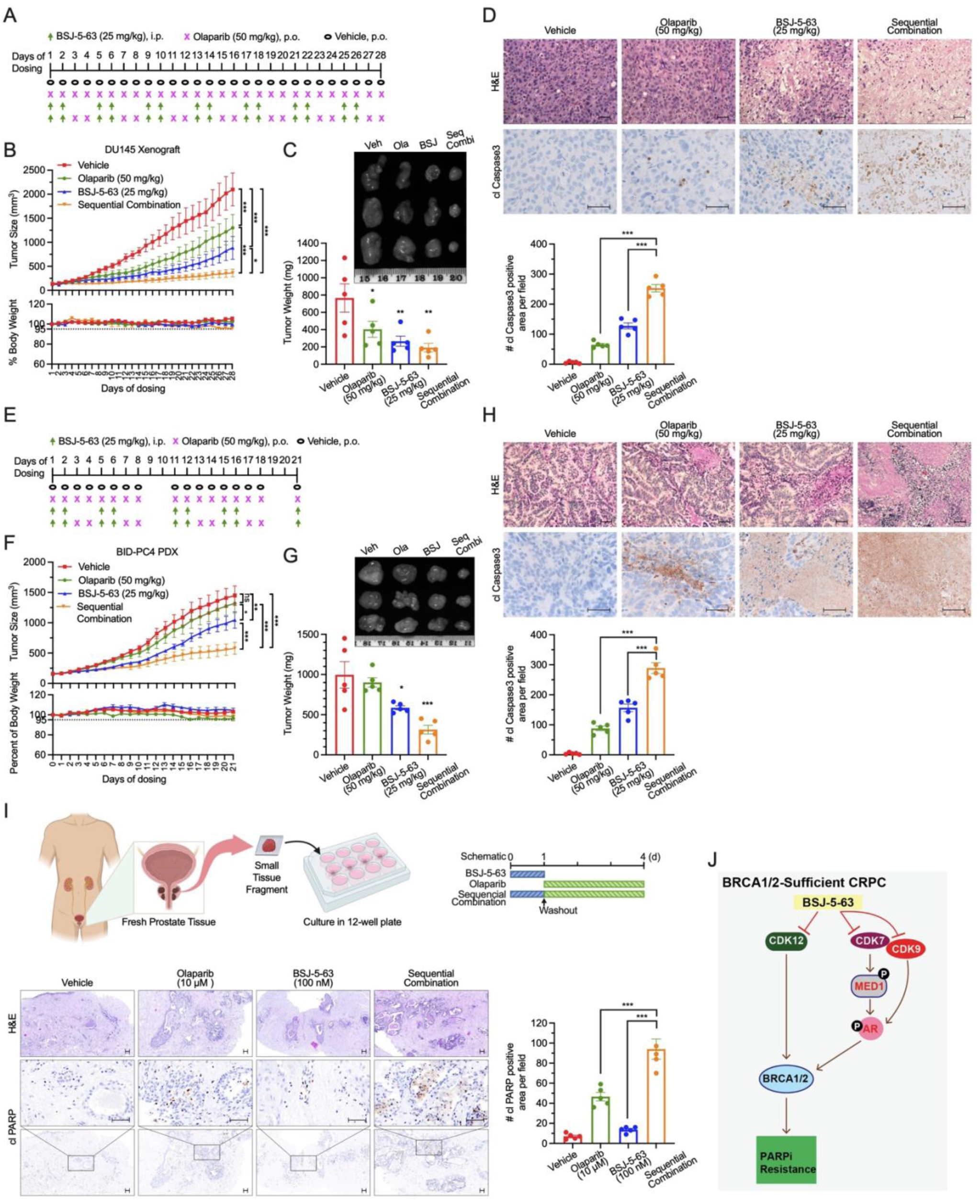
BSJ-5-63 sensitizes prostate tumors to PARP inhibition *in vivo.* (A) Schematic of sequential treatment protocol for DU145 xenograft. (B) The *in vivo* antitumor efficacy of sequential combination of BSJ-5-63 and olaparib in DU145 xenograft model. Top: Tumor volume curve. Bottom: Body weight curve of the mice. Data are presented as mean ± SEM (n = 5 per group). \**P* < 0.05, \*\*\**P* < 0.001; one-way ANOVA with Tukey’s multiple comparisons test. (C) Tumor weights of DU145 xenografts dissected on Day 28 of drug treatment with representative photos. Data are presented as mean ± SEM (n = 5 per group). \**P* < 0.05, \*\**P* < 0.01. (D) Immunohistochemistry analysis and H&E staining showing significantly increased apoptosis in DU145 tumors after sequential combination therapy. Scale bar: 50 μm. Right: Quantification of cl Caspase3 staining in the xenografts. Data were presented as mean ± SEM (n = 5 per group). \*\*\**P* < 0.001; one-way ANOVA with Tukey’s multiple comparisons test. (E) Schematic of sequential treatment protocol for BID-PC4 PDX model. (F) The *in vivo* antitumor efficacy of sequential combination of BSJ-5-63 and olaparib in BID-PC4 PDX model. Top: Tumor volume curve. Bottom: Body weight curve of the mice. Data are presented as mean ± SEM (n = 5 per group). *P < 0.05, \*\**P* < 0.01, \*\*\**P* < 0.001; one-way ANOVA with Tukey’s multiple comparisons test. (G) Tumor weights of BID-PC4 PDX dissected on Day 21 of drug treatment with representative photos. Data are presented as mean ± SEM (n = 5 per group). \**P* < 0.05, \*\*\**P* < 0.001. (H) Immunohistochemistry analysis and H&E staining showing significantly increased apoptosis in BID-PC4 PDX tumors after sequential combination therapy. Scale bar: 50 μm. Right: Quantification of cl Caspase3 staining in the PDX. Data are presented as mean ± SEM (n = 5 per group). \*\*\**P* < 0.001; one-way ANOVA with Tukey’s multiple comparisons test. (I) Short-term *ex vivo* cultures of PCa tissue fragments from patient #1 (Gleason score 7a / ISUP grade group 2). Top: Schematic diagram of the experimental procedure with sequential combination treatment protocol. Left Bottom: Immunohistochemistry analysis and H&E staining showing apoptosis levels in PCa tissues from short-term *ex vivo* cultures. Scale bar: 50 μm. Right Bottom: Quantification of cl-PARP staining. Data are presented as mean ± SEM. ***P < 0.001; one-way ANOVA with Tukey’s multiple comparisons test. (J) Model depicting the mechanism by which BSJ-5-63 induces acute BRCAness and AR signaling blockage through CDK12/7/9 degradation, enhancing PARPi sensitivity in PCa.

Furthermore, we used a newly developed BID-PC4 patient-derived xenograft (PDX) model, which lacks HRR deficiency. The key genomic alterations of BID-PC4 were determined by targeted exome sequencing (table S6). This model was insensitive to olaparib treatment, as evidenced by tumor growth in SCID mice. BSJ-5-63 demonstrated only a marginal inhibitory effect on tumor growth in this context (Fig. 7, E to G). Nevertheless, when the same sequential treatment approach was applied, combination therapy exerted a substantial inhibitory effect on tumor growth. Histological examination (H&E staining) revealed that the structure of PCa remained largely intact in the control and single agent-treated groups, whereas the combination therapy led to massive apoptotic cell death, as evidenced by cleaved caspase 3 staining (Fig. 7H). Finally, we extended our findings by using short-term *ex vivo* cultures of three patient-derived prostate tumors obtained from radical prostatectomy. We observed significantly increased apoptosis in PCa tissues after sequential combination treatment with BSJ-5-63 followed by olaparib as indicated, compared to single agent treatment, as evidenced by cleaved PARP1 staining (Fig. 7I and fig. S7).

## Discussion

In this study, we demonstrated the efficacy of BSJ-5-63 in degrading CDK12, CDK7, and CDK9, supporting its potential as a versatile therapeutic agent (Fig. 7J). The degradation of CDK12, coupled with the downregulation of long HRR genes, including BRCA1/2, aligns with previous studies highlighting the importance of CDK12 in controlling HRR function (*22, 45*). This degradation induces a state of BRCAness, rendering CRPC cells sensitive to PARP inhibition. The temporal persistence of this BRCAness state following BSJ-5-63 treatment paves the way for a rational sequential combination therapy, potentially minimizing toxicity and overcoming resistance issues commonly associated with single agent treatments. The extended mechanism of BSJ-5-63, targeting CDK7 and CDK9, further enhances its therapeutic potential. The inhibition of these kinases not only disrupts AR signaling, a primary therapeutic target in CRPC, but also interferes with the expression of AR-controlled HRR genes (*17, 18*), providing additional pathway to enhance BRCAness. RNA-seq analysis confirmed that the impaired HRR pathway is the primary mechanism of action induced by BSJ-5-63. Therefore, the proposed combinatorial approach involving the degradation of CDK12, CDK7, and CDK9 holds significant promise for a broad spectrum of CRPC patients, extending beyond those with BRCA1/2 alterations.

Both genetic deletion of CDK12 and suppression of its activity using BSJ-5-63 demonstrated that acute CDK12 inhibition can sensitize CRPC cells to PARP inhibition. However, prolonged CDK12 deletion in PCa cells led to a gradual and partial restoration of BRCA1/2 expression, resulting in a subsequent loss of sensitivity to PARP inhibition. This observation is in line with clinical data showing the insensitivity of PCa with CDK12 loss-of-function mutations (*24, 27–29*), indicating the complexity of cellular responses to CDK12 genetic alterations and prompting further exploration of the underlying mechanisms. These mechanisms may include substantial functional redundancy between CDK12 and its paralog kinase, CDK13 (*47, 48*). Furthermore, our results revealed that BSJ-5-63 can target AR-positive CRPC cells by degrading CDK7 and CDK9, key players in AR-mediated transcription. Targeting AR through CDK7 and CDK9, without involving AR ligand binding, may prove effective in CRPC with AR amplification, mutations or variants. Inhibition of AR signaling by BSJ-5-63 offers an alternative therapeutic strategy when direct AR-targeted therapies fail, potentially overcoming ARPI resistance, a major challenge in CRPC treatment. *In vivo* experiments using cell line-derived xenograft and PDX models demonstrated the translational potential of BSJ-5-63. While both olaparib and BSJ-5-63 as single agents showed limited impact, the sequential combination of BSJ-5-63 with olaparib produced synergistic effects, indicating the clinical relevance of this approach. Sequential combination therapy was not only well tolerated but also effective for CRPC, irrespective of genetic alteration status. *Ex vivo* experiments also demonstrated the efficacy of the sequential combination therapy, reinforcing its potential application in patients.

While both BSJ-5-63 and BSJ-4-116 degrade CDK12 and suppress HRR genes BRCA1/2, BSJ-5-63 distinguishes itself as a CDK12/7/9 triple degrader. This distinction is likely attributed to the use of the E3 ubiquitin ligase VHL ligand, as opposed to CRBN in BSJ-4-116. Consequently, BSJ-5-63 has an additional impact on AR signaling by targeting both CDK7 and CDK9, in contrast to BSJ-4-116, which targets only CDK9 in PCa cells. This differential targeting led to more potent inhibition of the AR pathway with BSJ-5-63, as CDK7 acts as a master regulator in AR-mediated transcription (*36*). Importantly, BSJ-5-63 exhibited favorable pharmacokinetic properties, including a longer half-life, reduced clearance rate, and a higher maximum plasma concentration than BSJ-4-116. These characteristics suggest that BSJ-5-63 may offer enhanced efficacy in vivo. However, our study has limitations. The potential off-target effects of BSJ-5-63, a common concern with compounds of this class, remain unexplored, which could affect its clinical utility and safety profile. Importantly, the primary toxicity may arise from targeting CDK7, CDK9, and CDK12, which are key transcriptional regulators. While these CDKs are well-established cancer therapy targets, with various inhibitors and degraders in development, further research is needed to determine if simultaneous inhibition of multiple CDKs is essential. The specific contributions of CDK7 and CDK9 degradation in inducing BRCAness require further validation.

In conclusion, clinical investigations over the past several years have shown that PARP inhibition is particularly effective in the context of BRCA1/2 alterations. However, the extent to which patients with non-BRCA1/2 genomic alterations can benefit from PARPis remains unclear, as gene-by-gene analyses have been inconclusive (*49, 50*). Indeed, mutations in non-BRCA HRR genes may not induce an HRR deficiency comparable to that caused by BRCA1/2 loss. Even in patients with BRCA mutations, acquired resistance can develop due to BRCA reversion mutations (*51*), which restore BRCA1/2 expression and HRR function. Therefore, it is critical to develop approaches that effectively suppress BRCA1/2 expression and synergize PARP inhibition. One such approach involves ARPIs, which have been shown to downregulate HRR gene expression to some extent in preclinical models (*43*). However, the clinical benefit of combining AR and PARP inhibition may not be attributable to AR-mediated transcription of the HRR genes (*21*). Further evidence is needed to determine whether ARPIs can induce a true BRCAness state. Our findings present a promising avenue for CRPC treatment through the distinctive profile of BSJ-5-63, which targets CDK12, CDK7, and CDK9 to transiently but thoroughly abolish BRCA1/2 expression and HRR function. Combination therapies with PARPis exhibit remarkable efficacy in preclinical models, laying the groundwork for future clinical studies.

## Materials and Methods

### Cells

Parental PCa cell lines were obtained from American Type Culture Collection (Manassas, VA, USA). LNCaP (Cat. CRL-1740, RRID: CVCL_0395), C4-2B (Cat. CRL-3315, RRID: CVCL_4784), 22Rv1 (Cat. CRL-2505, RRID: CVCL_1045), DU145 (Cat. HTB-81, RRID: CVCL_0105), and LAPC4 (Cat. CRL-13009, RRID: CVCL_4744) cells were cultured in RPMI 1640 medium (Gibco, Cat. 11875093). VCaP (Cat. CRL-2876, RRID: CVCL_2235) cells were grown in DMEM medium with Glutamax (Gibco, Cat. 10566016). HEK293T (Cat. CRL-3216, RRID: CVCL_0063) cells were grown in DMEM medium (Gibco, Cat. 21013024). The enzalutamide-resistant cell line MR49F (RRID: CVCL_RW53) was obtained from Dr. Amina Zoubeidi (University of British Columbia, Canada) and maintained in RPMI 1640 medium supplemented with 10 μM enzalutamide (*46*). The medium was supplemented with 10% fetal bovine serum (FBS) (Sigma, Cat. F0926) and 1% penicillin-streptomycin solution (Gibco, Cat. 15140122). The cell lines were maintained in an incubator at 37°C and 5% CO2. All cell lines were tested negative for mycoplasma contamination using MycoAlert (Lonza, Cat. LT07-318).

### Animals

Male SCID mice aged 4-5 weeks were obtained from Taconic Laboratories and acclimated for 1 week in a pathogen-free enclosure prior to the start of the study. Mice were maintained in a 12-hour light/12-hour dark cycles with free access to food and water. All procedures were conducted in compliance with the guidelines of the Institutional Animal Care and Use Committee (IACUC) at the Center for Comparative Medicine in Brigham and Women’s Hospital.

### Human prostate carcinoma tissue

Fresh prostate carcinoma tissue samples were collected at Goethe University Hospital Frankfurt, Germany from patients undergoing radical prostatectomy under license UCT-38-2023 approved by the UCT Scientific Board and Ethics Committee, and compliant with all relevant ethical regulations regarding research involving human participants. Signed consent was obtained from all patients. Samples were de-identified prior to transport to the laboratory.

### Compounds

BSJ-5-63, BSJ-5-63-NC, BSJ-4-116, and BSJ-4-116-NC were synthesized in-house. The medicinal chemistry and pharmacological profiles of the compounds are described in Supporting Information. Olaparib (Cat. HY-10162), rucaparib (Cat. HY-10617A), niraparib (Cat. HY-10619), talazoparib (Cat. HY-16106), and sunitinib (Cat. HY-10255A) were purchased from MedChem Express (Monmouth Junction, NJ, USA). For cell cultures, all compounds were dissolved in dimethyl sulfoxide (DMSO) (Sigma, Cat. D8418) to prepare a stock solution (10 mM) and stored at −20°C. The final dosing solution was prepared on the day of use by diluting the stock solution. For *in vivo* assays, the compounds were dissolved in a vehicle composed of 5% DMSO, 15% (w/v) Kolliphor HS 15 (Sigma, Cat. 42966), and 80% normal saline.

### Synthetic procedures

Unless otherwise noted, reagents and solvents were obtained from commercial suppliers and were used without further purification. 1H NMR spectra were recorded on 500 MHz Bruker Avance III spectrometer, and chemical shifts are reported in parts per million (ppm, δ) downfield from tetramethylsilane (TMS). Coupling constants (J) are reported in Hz. Spin multiplicities are described as s (singlet), br (broad singlet), d (doublet), t (triplet), q (quartet), and m (multiplet). Mass spectra were obtained on a Waters Acquity UPLC. Preparative HPLC was performed on a Waters Sunfire C18 column (19 mm × 50 mm, 5 μM) using a gradient of 15−95% methanol in water containing 0.05% trifluoroacetic acid (TFA) over 22 min (28 min run time) at a flow rate of 20 mL/min. Assayed compounds were isolated and tested as TFA salts. Purities of assayed compounds were in all cases greater than 95%, as determined by reverse-phase HPLC analysis.

### Synthesis of (2*S*,4*R*)-1-((*S*)-2-(11-((*R*)-3-((5-chloro-4-((2-(isopropylsulfonyl)phenyl)amino)pyrimidin-2-yl)amino)piperidin-1-yl)undecanamido)-3,3-dimethylbutanoyl)-4-hydroxy-*N*-((*S*)-1-(4-(4-methylthiazol-5-yl)phenyl)ethyl)pyrrolidine-2-carboxamide (BSJ-5-63)

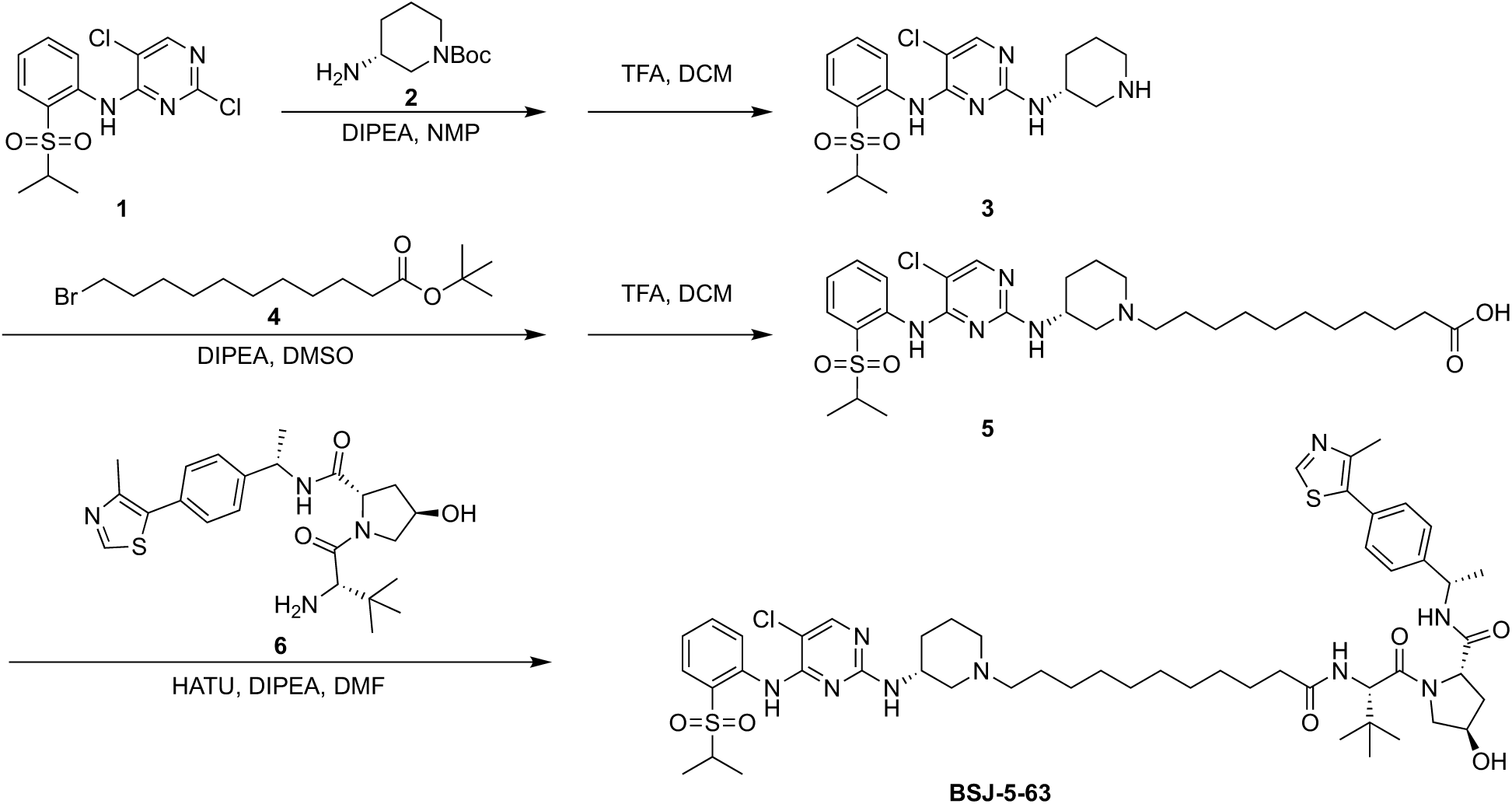

To a solution of 2,5-dichloro-*N*-(2-(isopropylsulfonyl)phenyl)pyrimidin-4-amine **1** (345 mg, 1.0 mmol) in 5 mL of NMP was added *tert*-butyl (*R*)-3-aminopiperidine-1-carboxylate (**2**) (300 mg, 1.5 mmol) and DIPEA (0.52 mL, 3.0 mmol). The reaction mixture was heated to 120°C and kept stirring overnight. The mixture was then warmed to room temperature, extracted with 100 mL of Ethyl Acetate (EA) and 50 mL of water. The organic layer was washed with 50 mL of Saturated Na_2_CO_3_ and 50 mL of brine, dried over anhydrous Na_2_SO_4_, and evaporated to give a yellowish residue which was directly dissolved into 2.5 mL of DCM, followed by slow addition of 2.5 mL of TFA at ice-bath. The mixture was warmed to room temperature and stirred for 0.5h. The solvent was then evaporated, and the residue was purified by reverse phase HPLC (5-95% MeOH in H_2_O) to give **3** (TFA salt) as a yellow solid (368 mg, 90% in two steps). LC-MS: m/z 410 (M+1).

To a solution of **3** (90 mg, 0.22 mmol) in DMSO (2 mL) was added *tert*-butyl 11-bromoundecanoate (**4**, 130 mg, 0.44 mmol) and DIPEA (0.115 mL, 0.66 mmol). The mixture was heated to 80°C and kept stirring for 24h. The mixture was then cooled down to room temperature, extracted, dried, filtered and concentrated to give a light brown residue which was then dissolved into 2 mL of DCM, followed by slow addition of 1 mL TFA at O°C. The mixture was then stirred at room temperature for 1h, and evaporated. The residue was purified by reverse phase HPLC (5-95% MeOH in H_2_O) to give **5** as a light grey solid (59 mg, 45% in two steps). LC-MS: m/z 594 [M+1].

To a solution of **5** (25 mg, 0.0422 mmol) in 2 mL of DMF was added VHL ligand (2*S*,4*R*)-1- ((*S*)-2-amino-3,3-dimethylbutanoyl)-4-hydroxy-*N*-((*S*)-1-(4-(4-methylthiazol-5-yl)phenyl)ethyl)pyrrolidine-2-carboxamide (**6**,19 mg, 0.0422 mmol), HATU (33 mg, 0.0844 mmol) and DIPEA (37 μL, 0.211 mmol). The resulting mixture was stirred for 1h at room temperature, then evaporated the solvent and purified by reverse phase HPLC (5-95% MeOH in H_2_O) to give **BSJ-5-63** (TFA salt) as an off-white solid (36 mg, 83%). LC-MS: *m/z* 1021 [M+1]. ^1^H NMR (500 MHz, DMSO-*d*_6_) δ 9.53 (d, *J* = 5.4 Hz, 1H), 9.39 (br, 1H), 8.98 (s, 1H), 8.36 (d, *J* = 7.8 Hz, 1H), 8.17 (s, 1H), 7.88-7.81 (m, 1H), 7.77 (d, *J* = 9.1 Hz, 2H), 7.45-7.40 (m, 2H), 7.39-7.32 (m, 4H), 4.95-4.86 (m, 1H), 4.50 (d, *J* = 9.3 Hz, 1H), 4.41 (t, *J* = 8.1 Hz, 1H), 4.31-4.25 (m, 1H), 4.11 (br, 1H), 3.31 (br, 6H), 3.06 (br, 2H), 2.87-2.73 (m, 1H), 2.45 (s, 3H), 2.28-2.20 (m, 1H), 2.13-2.05 (m, 2H), 2.04-1.89 (m, 2H), 1.83-1.75 (m, 1H), 1.53-1.41 (m, 4H), 1.37 (d, *J* = 7.0 Hz, 3H), 1.24 (d, *J* = 12.2 Hz, 14H), 1.17 (t, *J* = 6.2 Hz, 6H), 0.92 (s, 9H). ^13^C NMR (126 MHz, DMSO) δ 172.62, 171.10, 170.07, 152.01, 148.17, 145.09, 135.55, 131.61, 131.51, 130.14, 129.29, 126.85, 69.22, 59.01, 56.83, 56.69, 55.46, 48.18, 38.18, 35.67, 35.38, 29.27, 29.19, 29.12, 28.89, 26.93, 26.88, 26.73, 26.41, 25.88, 23.70, 22.84, 16.40, 15.36, 15.30, 15.25.

### Synthesis of (2*R*,4*S*)-1-((*S*)-2-(11-((*R*)-3-((5-chloro-4-((2-(isopropylsulfonyl)phenyl)amino)pyrimidin-2-yl)amino)piperidin-1-yl)undecanamido)-3,3-dimethylbutanoyl)-4-hydroxy-*N*-((*S*)-1-(4-(4-methylthiazol-5-yl)phenyl)ethyl)pyrrolidine-2-carboxamide (BSJ-5-63-NC)

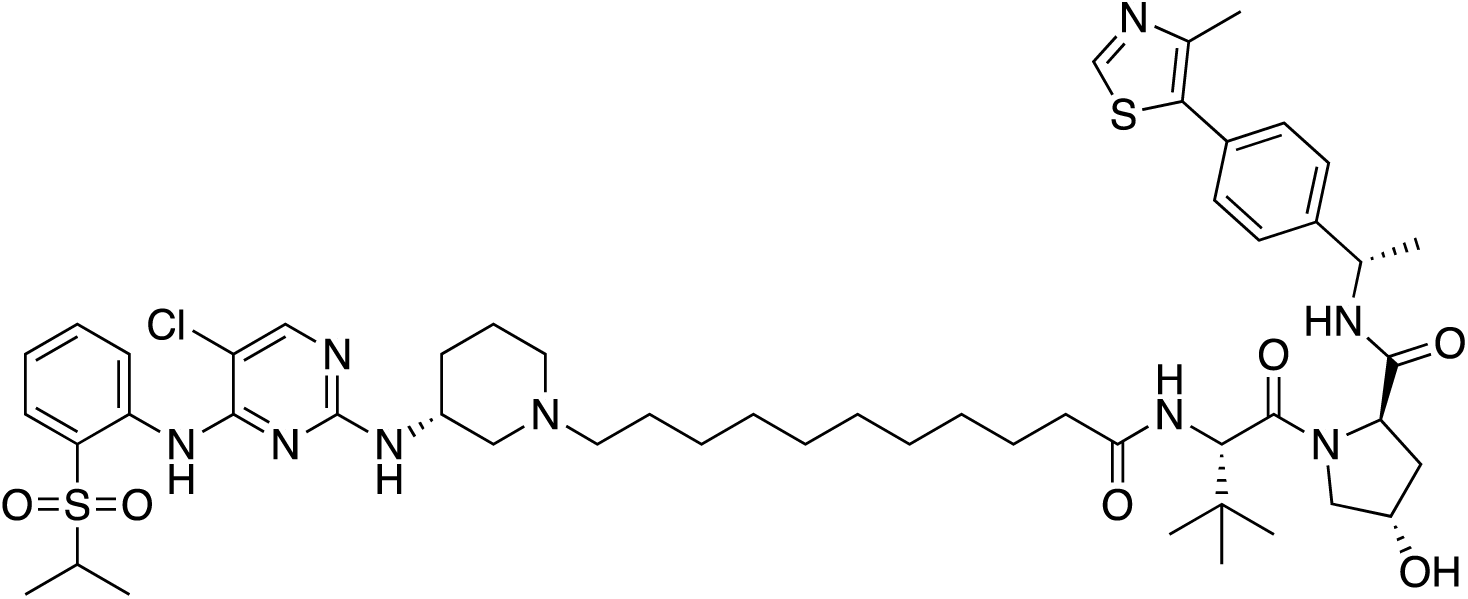

**BSJ-5-63-NC** was synthesized with similar procedures as **BSJ-5-63** from **5** and (2*R*,4*S*)-1- ((*S*)-2-amino-3,3-dimethylbutanoyl)-4-hydroxy-*N*-((*S*)-1-(4-(4-methylthiazol-5-yl)phenyl)ethyl)pyrrolidine-2-carboxamide. **BSJ-5-63-NC** (TFA salt) was obtained as an off-white solid. LC-MS: *m/z* 1021 [M+1]. ^1^H NMR (500 MHz, DMSO-*d*_6_) δ 9.48 (s, 1H), 8.97 (d, *J* = 6.1 Hz, 1H), 8.74 (br, 1H), 8.13 (br, 1H), 8.02 (d, *J* = 8.1 Hz, 1H), 7.94-7.87 (m, 1H), 7.83 (dd, *J* = 8.0, 1.6 Hz, 1H), 7.73 (s, 1H), 7.49-7.41 (m, 4H), 7.32 (t, *J* = 7.7 Hz, 1H), 5.13 (d, *J* = 4.1 Hz, 1H), 4.97-4.85 (m, 1H), 4.43-4.34 (m, 2H), 4.34-4.26 (m, 1H), 3.82 (dd, *J* = 10.4, 5.4 Hz, 1H), 3.50 (dd, *J* = 10.5, 4.3 Hz, 1H), 3.47-3.39 (m, 1H), 3.34 (br, 6H), 3.18 (d, *J* = 5.0 Hz, 1H), 2.46 (s, 3H), 2.32-2.14 (m, 2H), 2.11-2.02 (m, 1H), 2.02-1.91 (m, 2H), 1.84 (br, 2H), 1.58-1.36 (m, 4H), 1.31 (d, *J* = 7.0 Hz, 3H), 1.26-1.08 (m, 20H), 0.97 (s, 9H). ^13^C NMR (126 MHz, DMSO) δ 173.38, 171.00, 170.17, 155.13, 151.89, 148.19, 144.89, 138.86, 131.41, 130.12, 129.24, 127.24, 127.02, 123.18, 68.84, 59.15, 57.56, 56.50, 55.67, 55.64, 55.42, 55.39, 47.96, 38.25, 35.23, 34.70, 29.42, 29.38, 29.30, 29.21, 29.18, 29.08, 26.93, 26.81, 25.78, 22.99, 19.04, 16.47, 15.34, 15.32.

### Generation of gene KO cell lines using CRISPR/Cas9

LentiGuide-Puro (Addgene, Cat. 52963, RRID: Addgene_52963) and lentiCas9-Blast (Addgene, Cat. 52962, RRID: Addgene_52962) plasmids were used to generate CDK12 and CDK9 KO cell lines. CRISPR sgRNAs targeting CDK12 and CDK9 were cloned into a lentiGuide-Puro vector. AAVS1 sgRNAs were cloned into the same vector and served as controls. The sgRNAs used in this study are listed in table S6. Lentiviruses were generated by transfecting sub-confluent HEK293T cells along with the lentiviral vector and packaging vectors pCMV-VSV-G (Addgene, Cat. 8454, RRID: Addgene_8454) and psPAX2 (Addgene, Cat. 12260, RRID: Addgene_12260) at a 2:1:1 ratio using Lipofectamine 3000 transfection reagents (Invitrogen, Cat. L3000015). Viral supernatants were collected 48 hours after transfection and filtered through a 0.45-μm membrane (Corning, Cat. 431220). For cell infection, 1.5 mL of viral supernatant, 0.5 mL fresh medium and 8 μg·mL^−1^ polybrene were added to 5 million cells per well in six-well plates. Plates were centrifuged at 600x g for 90 min at 37°C during the spin infection. The virus-containing medium was removed after 24 hours. Initially, cells overexpressing Cas9 were generated under 20 μg·mL^−1^ Blasticidin selection. Subsequently, CDK12 and CDK9 KO cells were further generated under 2 μg·mL^−1^ puromycin selection. The KO efficiency was validated using Western blotting.

### Western blotting

Cells were washed with PBS buffer (Gibco, Cat. 10010023) and lysed in 1xRIPA lysis buffer (Thermo Scientific, Cat. 89901) supplemented with protease and phosphatase inhibitor cocktail (Thermo Fisher, Cat. 78442). The total protein concentration was measured using Pierce BCA Protein Assay Kit (Thermo Fisher, Cat. 23227). Western blotting was performed as previously described (*52*). Briefly, Equal amounts of proteins were separated by SDS-PAGE and transferred to PVDF membranes (Millipore, Cat. IPVH00010). Membranes were incubated with primary antibodies in 5% BSA in 1x TBST buffer overnight at 4°C. After primary antibody incubation, membranes were probed with a secondary antibody at room temperature for 1 hour. The antibodies used in this study are listed in table S6.

### Immunofluorescence staining

CDK12-KO 22Rv1 and AAVS1 control cells were seeded into four-well Millicell EZ slide (Millipore, Cat. PEZGS0416) pre-coated with Poly-L-Lysine (Sigma, Cat. P4707-50ml) overnight. Cells were treated with 100 nM BSJ-5-63 for 24 hours before irradiation (8 Gy). After treatment, the cells were washed three times with PBS and then fixed in 4% formaldehyde (Electron Microscopy Sciences, Cat. 15714) for 10 minutes at room temperature. Fixed cells were further incubated with 0.1%Triton X-100 in PBS for 10 minutes to permeabilize the cells and then washed with PBST (0.1% Tween20 in PBS) three times for 5 minutes. Fixed cells were blocked in PBS containing 5% BSA for H2AX or in PBS containing 1% BSA and 20% FBS for RAD51 at room temperature for 1 hour. After washing with PBST three times for 5 minutes each, cells were incubated with phospho-histone H2AX Ser139 or RAD51 antibodies diluted in the same blocking buffer overnight at 4°C. After washing with PBST, the cells were further incubated with the appropriate secondary antibodies for 1 hour at room temperature. Cells were washed with PBST, mounted, and counterstained with DAPI (Mounting Medium With DAPI-Aqueous, Fluoroshield, Abcam, Cat. ab104139) overnight at 4°C. The antibodies used are listed in table S6. Each experiment was performed independently in triplicate. To calculate the γ-H2AX foci, the number of foci was counted from at least 50 cells per replicate for each condition. To calculate RAD51 foci, the number of cells containing at least 5 foci was counted in 10 fields for each replicate under each condition.

### Colony formation assay

Cells were seeded at a density of 15,000 cells per well for LNCaP, C4-2B and DU145 cells, and 30,000 cells per well for 22Rv1 cells in six-well plates and incubated overnight before treatment as indicated. For the colony formation assay, cells were incubated for 14 days at 37°C with 5% CO_2_, with the medium changed once per week. For sequential combination treatment, cells were treated with BSJ-5-63 or BSJ-4-116 at the indicated dose for 2 days. Cells were rinsed with 1xPBS and then treated with PARPis olaparib, rucaparib, or niraparib at the indicated doses for an additional 12 days. The colonies were subsequently washed with PBS and stained with 0.1% Crystal Violet as previously described (*53*). Colony images were quantified using the ImageJ software (National Institutes of Health). Three independent experiments were performed.

### Cell viability assay

Cells were seeded at a density of 7,200 cells per well (for 3 days of treatment) or 2,700 cells per well (for 7 days of treatment) in 96-well plates and incubated overnight before treatment. Subsequently, the cells were treated with the indicated compounds. Cell viability was determined using alamarBlue Cell Viability Reagent (ThermoFisher, Cat. DAL1025) according to the manufacturer’s instructions. IC_50_ values were estimated using GraphPad Prism 9.0 (GraphPad Software). Three independent experiments were performed in triplicates.

### RNA Interference

For knockdown experiments, cells were transfected with 100 nM ON-TARGETplus SMARTpool siRNA targeting CDK7 (Dharmacon, L-003241-00-0005) or a non-targeting pool (Dharmacon, D-001810-10-05) as a negative control, using Lipofectamine RNAiMAX (Invitrogen, Cat. 13778150) according to the manufacturer’s instructions. Cells were harvested at 48 hours post-transfection.

### DHT Stimulation

LNCaP, CDK12-KO 22Rv1 and CDK9-KO 22Rv1cells were grown in medium containing charcoal stripped fetal bovine serum (CSS) for 3 days. Subsequently, the cells were treated with 10 nM 5-α-dihydrotestosterone (DHT, Sigma, Cat. D-073-1ML) in ethanol or with ethanol alone as a control. Cells were harvested at 16 hours post-stimulation. For DHT stimulation with BSJ-5-63, LNCaP cells grown in CSS-containing medium for 3 days were treated with BSJ-5-63 as indicated. Three hours post-treatment, the cells were stimulated with 10 nM DHT for an additional 13 hours. For DHT stimulation with CDK7 RNA interference, LNCaP cells grown in CSS-containing medium for 3 days were transfected with 100 nM ON-TARGETplus SMARTpool siRNA targeting CDK7 or a non-targeting pool as indicated. Thirty-two hours post-transfection, the cells were stimulated with 10 nM DHT for an additional 16 hours.

### RNA isolation and RT-qPCR

Total RNA was extracted using RNeasy Plus Mini Kit (Qiagen, Cat. 74134) according to the manufacturer’s protocol. One microgram of RNA was reverse transcribed to complementary DNA in 20 μL reaction volumes using iScript Reverse Transcription Supermix (Bio-Rad, Cat. 1708841) according to the manufacturer’s protocol. RT-qPCR was performed using SYBR Select Master Mix for CFX (Applied Biosystems, Cat. 4472942) using the Bio-Rad CFX Connect Real-Time System. Briefly, 100 ng of cDNA was used as a template in 20 μL reaction volumes with 250 nM primers. Amplification reactions were run for 40 thermocycles of 15 seconds at 95°C, 1 minute at 59°C. The primers used are listed in table S6.

### RNA-seq assay

22Rv1 cells were treated with DMSO, BSJ-5-63 (100 nM), or BSJ-5-63-NC (100 nM) for 8 hours. Treatments were performed in duplicated. Cells were then rinsed with ice-cold 1xPBS. Total RNA was extracted using the RNeasy Plus Mini Kit according to the manufacturer’s protocol. The samples were sent to Genewiz for RNA sequencing. RNA-seq data analysis was performed as previously described (*53*). RNA-seq data in this study have been deposited in the GEO database under the accession number GSE263218 and are publicly available.

### Proteomics

22Rv1 cells were seeded in 10 mm dishes overnight before treatment with DMSO, BSJ-5-63 (100 nM), BSJ-5-63-NC (250 nM), or BSJ-4-116 (100 nM) for 8 hours. Cells were rinsed with cold PBS and stored at −80°C. Treatments were performed in biological triplicate. Proteomics and data analysis were performed as previously described (*34*). The mass spectrometry proteomics data in this study are provided within tables S2 and S3.

### *In vitro* mouse liver microsomal stability assay

The stability of BSJ-5-63 and BSJ-4-116 was determined in mouse liver microsomes using the service of Scripps Florida (FL, USA). Mouse liver microsomes (0.5 mg/mL) were incubated with BSJ-5-63 (1μM) or BSJ-4-116 (1μM) at 37°C in the presence of the cofactor NADPH (1mM). Sunitinib, a multi-targetd receptor kinase inhibitor, was used as a positive control in parallel. Samples were collected at time points 0, 5, 10, 20, 40, and 60 minutes for the measurement of remaining compounds by LC-MS/MS.

### *In vivo* xenograft model

Cells were filtered through 70 μm cell strainers (BD, NJ, USA) and suspended in serum-free media. Xenografts were initiated by subcutaneous injection of cells (5 x 10^6^ cells/50μL/mouse with an additional 50μL Matrigel, n = 5 per group) into the right flank near the axillary fossa of the mice. Tumor growth was monitored daily using the standard formula: V = length×width^2^×0.5. When the tumor volume approached approximately 150 mm^3^, the mice were randomly assigned into treatment groups.

For BSJ-5-63 single reagent treatment, mice bearing 22Rv1 xenografts were randomly assigned to three groups (n = 5 per group) and treated with vehicle (5% DMSO/15% Solutol HS 15/80% saline, i.p., daily), BSJ-5-63 (50 mg/kg, i.p., daily), or BSJ-5-63 (50 mg/kg, i.p., 1 day on/2 days off). Mice were euthanized after 12 days of treatment. For sequential combination treatment, mice bearing 22Rv1 or DU145 xenografts were randomly assigned to four groups (n = 5 per group) and treated with vehicle (p.o.), olaparib (50 mg/kg, p.o.), BSJ-5-63 (25 mg/kg, i.p.), or a sequential combination of BSJ-5-63 with olaparib as indicated. Mice were euthanized after three or four weeks of treatment, or when the tumor length reached 2 cm. For olaparib single reagent treatment, mice bearing CDK12-KO 22Rv1 xenografts (using gRNA sg1 and sg2) and AAVS1-KO control xenografts were randomly assigned to two groups separately and treated with vehicle (p.o.) or olaparib (50 mg/kg, p.o.) daily. Mice were euthanized after three weeks of treatment, or when the tumor length reached 2 cm. All compounds were formulated in the vehicle. The injection volume was 0.1 mL/10 g of mice body weight. Daily body weight was monitored. Mice were euthanized three hours post-final dose, and tumors were excised, weighted, and subjected to routine histopathological examination.

### *In vivo* PDX model

The human PCa BID-PC4 PDX was generated through the Dana-Farber/Harvard Cancer Center Genitourinary Oncology Rapid Autopsy Program, as previously described (*54*). The PDX was subjected to targeted exome sequencing, and genetic alterations in key genes are listed in table S7. No deleterious biallelic mutations were identified in the HRR genes. BID-PC4 was maintained by constant passaging in male SCID mice. Intact male SCID mice were implanted subcutaneously with BID-PC4 tumor bits in 50μL of 1xPBS per mouse with an additional 50μL Matrigel into the right flank near the axillary fossa of mice. When tumors reached approximately 150 mm^3^, mice were randomly assigned to four groups (n = 5 per group) and treated with vehicle (5% DMSO/15% Solutol HS 15/80% saline, p.o.), olaparib (50 mg/kg, p.o.), BSJ-5-63 (25 mg/kg, i.p.), and sequential combination of BSJ-5-63 with olaparib as indicated. All compounds were formulated in the vehicle. The injection volume was 0.1 mL/10 g of mouse body weight. Daily body weight was monitored. Mice were euthanized after three weeks of treatment or when tumor length reached 2 cm. The tumors were excised, weighted, and subjected to routine histopathological examination.

### Haematoxylin and Eosin (H&E) staining

Tumor tissues were fixed in 4% formalin, embedded in paraffin, cut into 4-μm thick sections, and mounted onto slides. Standard staining with H&E was performed.

### Immunohistochemistry

Immunohistochemistry staining of 4-μm thick tumor tissue sections was conducted as previously described (*55*). Briefly, after dewaxing, rehydration, and antigen retrieval, the sections were blocked with 5% serum for 1 hour. Subsequently, the sections were incubated with primary antibodies overnight at 4°C, followed by incubation with Polymer Helper for 20 minutes and secondary antibody for another 20 minutes. After the color reaction, the slides were counterstained with haematoxylin.

### *In vivo* pharmacokinetic analysis

The *in vivo* pharmacokinetic analysis was conducted using the service of Scripps Florida (FL, USA). The pharmacokinetic properties of BSJ-5-63 and BSJ-4-116 were determined in male C57BL/6 mice following intraperitoneal administration of 10 mg/kg. The compounds were formulated in 5% DMSO/15% Solutol HS 15/80% saline. Blood samples were collected up to 8 hours post-dosing for the measurement of compound concentrations in the blood using LC-MS/MS.

### *In vivo* pharmacodynamic analysis

The *in vivo* pharmacodynamic properties and efficacy of BSJ-5-63 were determined using 22Rv1 xenografts in male SCID mice. Xenografts were initiated by subcutaneous injection of 22Rv1 cells (5 x 10^6^ cells/50μL/mouse with an additional 50μL Matrigel, n = 3 per group) into the right flank near the axillary fossa of the mice. Tumor growth was monitored daily using the standard formula: V = length×width^2^×0.5. Mice bearing tumors of approximately 200 mm^3^ were randomly assigned to three groups and treated with vehicle (5% DMSO/15% Solutol HS 15/80% Saline, i.p., bid), BSJ-5-63 (25 mg/kg, i.p., bid), or BSJ-5-63 (50 mg/kg, i.p., bid) for three days. Three hours after the final dose, the mice were euthanized, and the tumors were excised and subjected to Western blotting.

### *Ex vivo* human PCa patient model

Fresh prostate carcinoma tissue samples were collected at Goethe University Hospital Frankfurt, Germany from patients undergoing radical prostatectomy under license UCT-38-2023 approved by the UCT Scientific Board and Ethics Committee, and compliant with all relevant ethical regulations regarding research involving human participants. Signed consent was obtained from all patients. Samples were de-identified prior to transport to the laboratory. Tissue specimens from patients #1 (Gleason score 7a / ISUP grade group 2), #2 (Gleason Score 7a / ISUP grade group 2), and #3 (Gleason score 9 / ISUP grade group 5) were obtained from radical prostatectomy without prior therapy. Tissues were cut into small fragments and transferred into 12-well plates with 2 ml of culture medium per well (DMEM/F12 containing 10% FBS, 1% GlutaMAX, and 1% penicillin-streptomycin solution). The tissue slices were treated with 100 nM BSJ-5-63 for 24 hours followed by washout of BSJ-5-63 and treatment with 10 µM olaparib for 3-4 days. As controls, treatments with DMSO, BSJ-5-63 (100 nM), or olaparib (10 µM) were performed for 4-5 days as indicated. Multiple tumor pieces were used for each condition. The plates were incubated at 37°C and 5% CO2. Subsequently, tissue fragments were fixed in 4% paraformaldehyde for 24 hours and embedded in paraffin. Immunohistochemistry staining for cleaved PARP (Asp214) (Cat# 5625, Cell Signaling, dilution ratio 1:100) was performed, followed by counterstaining with haematoxylin.

### Statistical analysis

All data are presented as mean ± SEM. Comparisons between two groups were analyzed using two-tailed unpaired Student’s t-test. For multiple group comparisons, one-way or two-way ANOVA followed by the Tukey-Kramer test for post-hoc analysis was applied. Statistical analyses were performed using GraphPad Prism 9 (GraphPad software). The significance levels are denoted as * P < 0.05, ** P < 0.01, and *** P < 0.001.

## Acknowledgments

This work was supported in part by NIH R01CA262524, R01CA279410, R21CA267496, DoD W81XWH-22-1-0477 to L.J., and the DF/HCC Incubator Award to S.P.B. and L.J. We thank Drs. Nathanael Gray and Tinghu Zhang for providing the compounds BSJ-4-116 and BSJ-5-63, reviewing the manuscript, and engaging in insightful discussions. The development of BSJ-5-63 was supported in part by R01CA218278 to N.G., and the proteomics work was supported in part by the Cancer Prevention and Research Institute of Texas, Grant RR220032 to M. K. who is a CPRIT Scholar in Cancer Research.

## Author contributions

Conceptualization: FG, BJ, LJ

Methodology: FG, BJ, MK, AF, SPB, LJ

Investigation: FG, BJ, JJ, ZH, TTsujino, TTakai, SA, CP, JK, GAB, RE, MK, AF, SPB, ASK, LJ

Visualization: FG, BJ, JJ, CP, JK, GAB, RE, MK, AF, LJ

Supervision: LJ

Writing—original draft: FG, LJ

Writing—review & editing: FG, BJ, MK, AF, SPB, LJ

## Data and materials availability

All data needed to evaluate the conclusions in the paper are present in the main text or the Supplementary Materials.

## Supplementary Figures

**Fig. S1.**
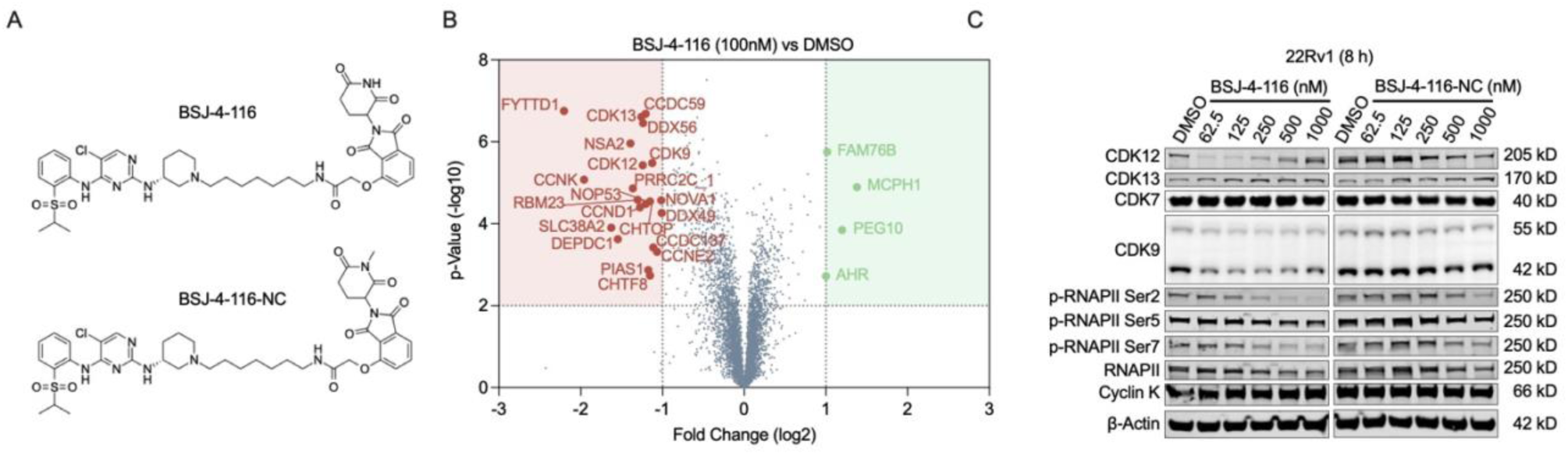
Characterization of CDK12 degrader BSJ-4-116. (A) Chemical structures of BSJ-4-116 and its negative control analog BSJ-4-116-NC. (B) Proteome-wide selectivity of BSJ-4-116. Quantitative proteomics showing the relative abundance of proteins measured by multiplexed quantitative MS-based proteomics in 22Rv1 cells treated with BSJ-4-116 (100 nM) or DMSO for 8 hours. (C) Western blots of indicated proteins in 22Rv1 cells after 8 hours of treatment with BSJ-4-116 or BSJ-4-116-NC in a dose-dependent manner. Representative blots from three independent experiments are shown.

**Fig. S2.**
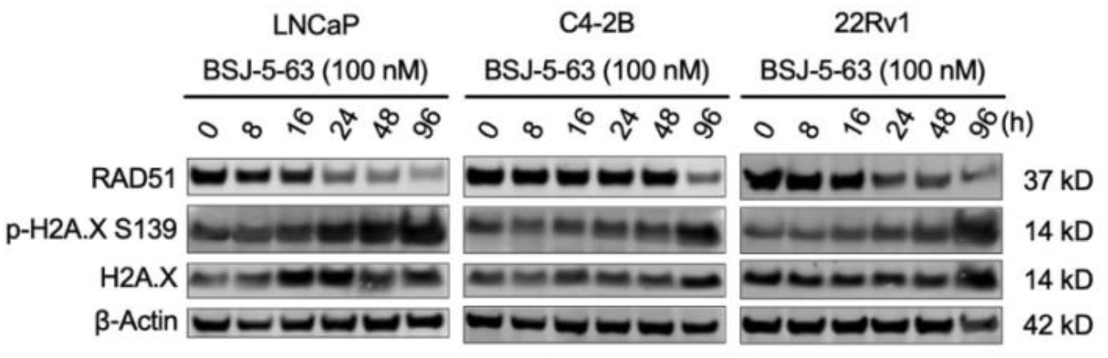
CDK12 degradation by BSJ-5-63 reduces RAD51 expression and induces DNA damage. Western blots of indicated proteins in LNCaP, C4-2B and 22Rv1 cells after BSJ-5-63 treatment in a time-course manner. Representative blots from three independent experiments are shown.

**Fig. S3.**
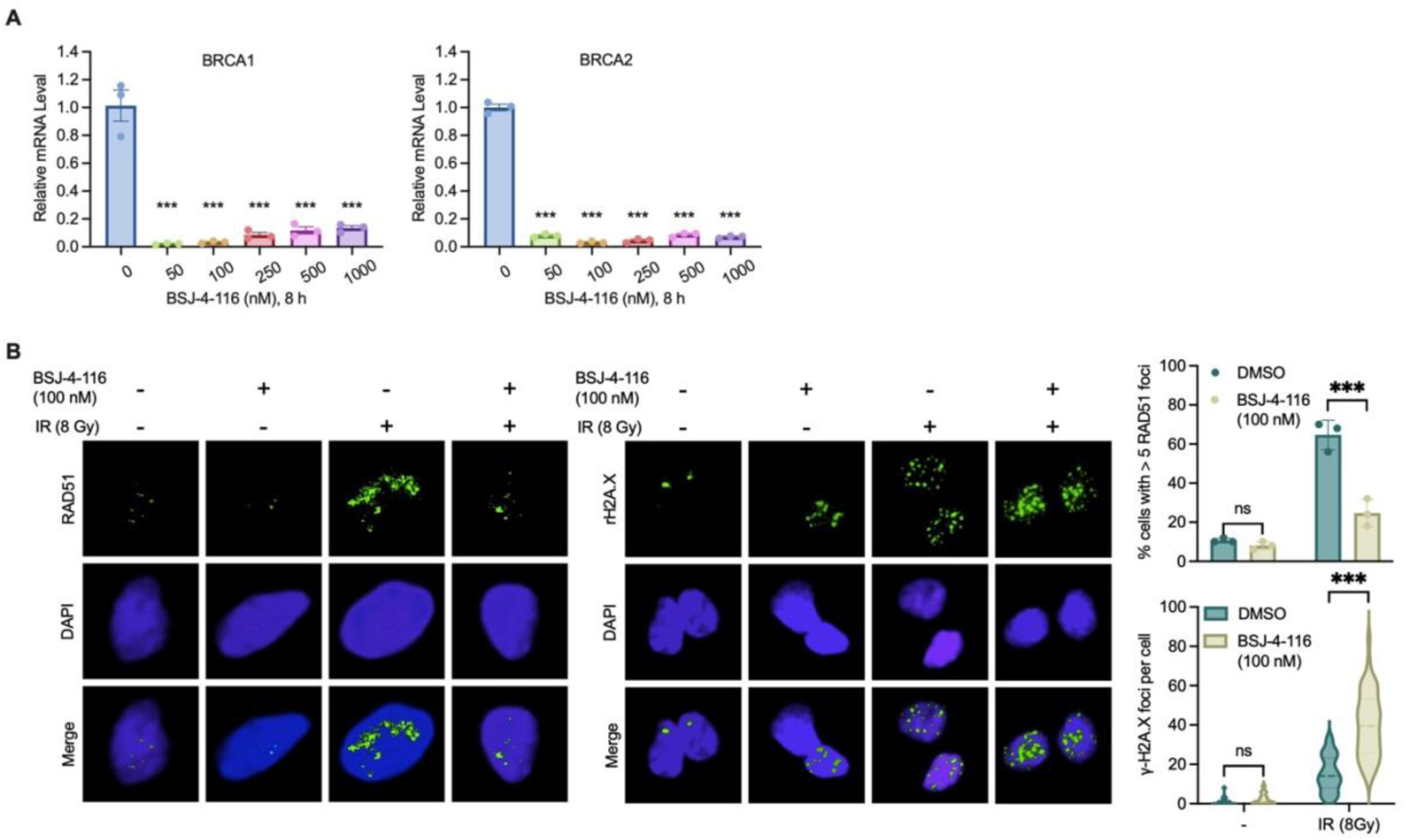
CDK12 degradation by BSJ-4-116 induces acute BRACness and impairs HRR. (A) mRNA levels of BRCA1/2 were measured by real-time RT-PCR. Data are presented as mean ± SEM (n=3). \*\*\**P* < 0.001; one-way ANOVA with Tukey’s multiple comparisons test. (B) Representative images of immunofluorescence staining and quantification of the number of RAD51 and γ-H2AX foci in 22Rv1 cells 24 hours after BSJ-4-116 (100 nM) treatment followed by 8 Gy irradiation. Representative images from three independent experiments. \*\*\**P* < 0.001; foci were counted in at least 50 cells for each replicate under each condition (n =3). Two-way ANOVA with Tukey’s multiple comparisons test.

**Fig. S4.**
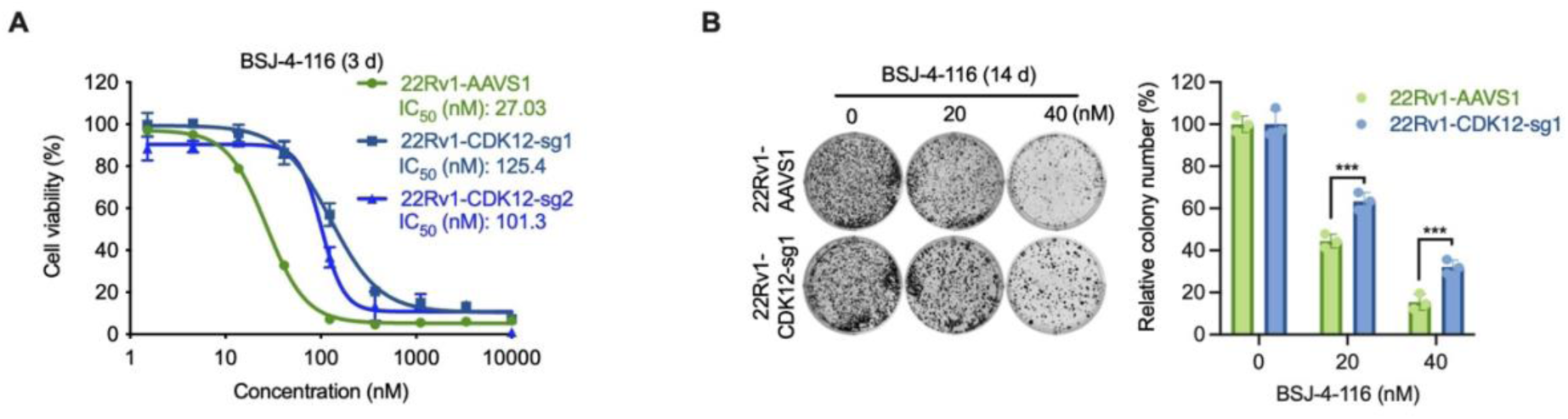
CDK12 KO reduces the antiproliferative effect of BSJ-4-116. (A) Cell viability assay showing reduced antiproliferative effect of BSJ-4-116 in CDK12-KO 22Rv1 cells. Data are presented as mean ± SEM (n = 3). \*\*\**P* < 0.001; one-way ANOVA with Tukey’s multiple comparisons test. (B) Colony formation assay showing reduced antiproliferative effect of BSJ-4-116 in CDK12-KO 22Rv1 cells. Representative colony images from three independent experiments are shown. Data are presented as mean ± SEM (n = 3). \*\*\**P* < 0.001; one-way ANOVA with Tukey’s multiple comparisons test.

**Fig. S5.**
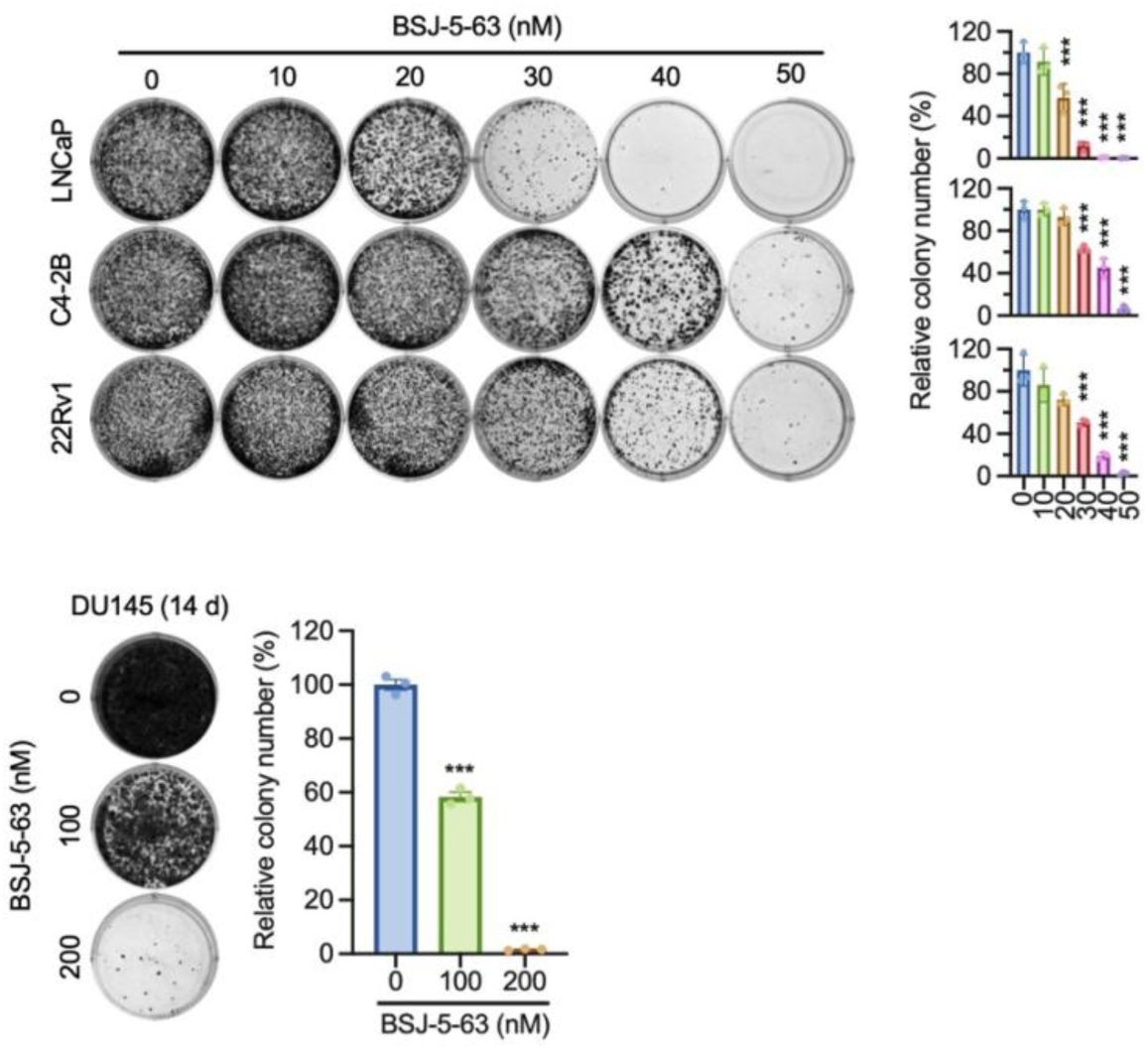
Inhibition of colony formation by BSJ-5-63. LNCaP, C4-2B, 22Rv1, and DU145 cells were treated with a dose range of BSJ-5-63 or DMSO for 14 days. Representative colony images from three independent experiments are shown. Data are presented as mean ± SEM (n = 3). \*\*\**P* < 0.001; one-way ANOVA with Tukey’s multiple comparisons test.

**Fig. S6.**
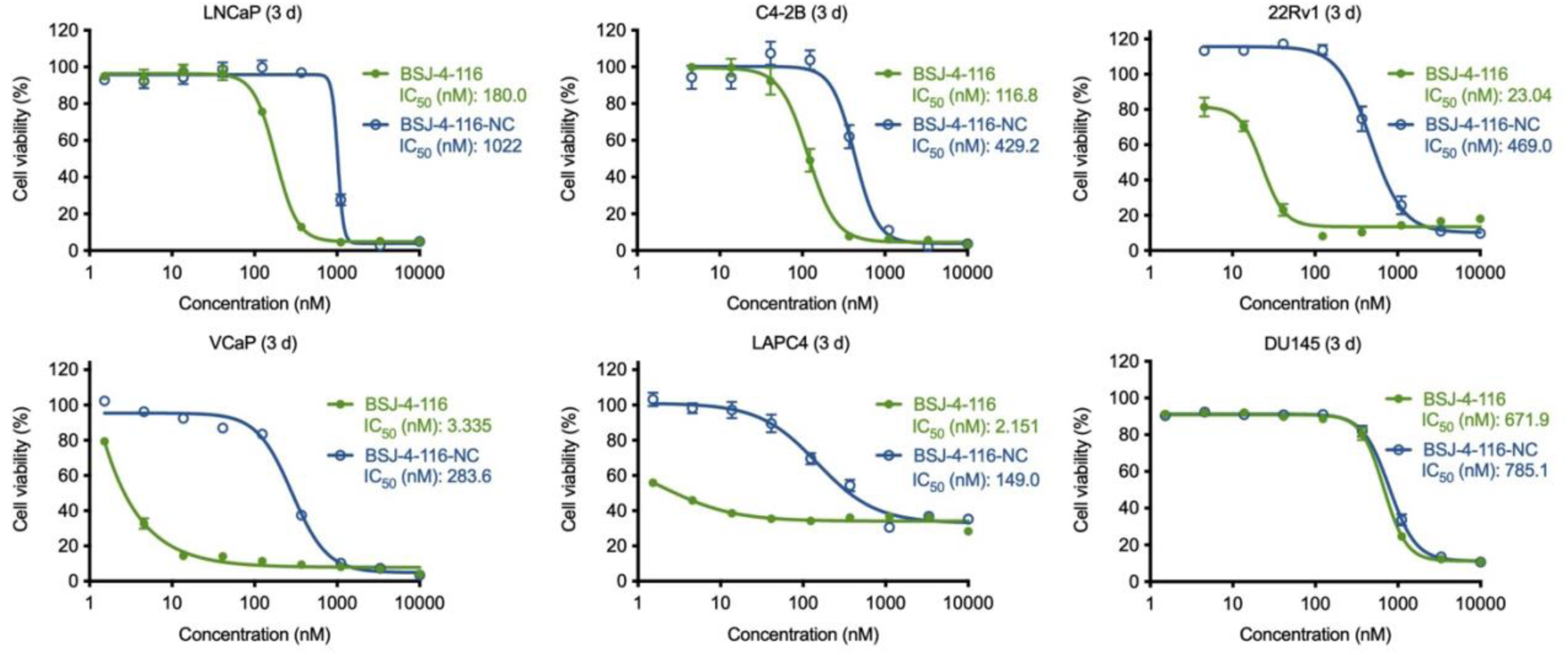
CDK12 degrader BSJ-4-116 inhibits PCa cell growth. Dose-response curves for various human PCa cells treated with BSJ-4-116 or BSJ-4-116-NC at the indicated dose range for 72 hours. IC_50_ was calculated and shown. Data are presented as mean ± SEM (n = 3).

**Fig. S7.**
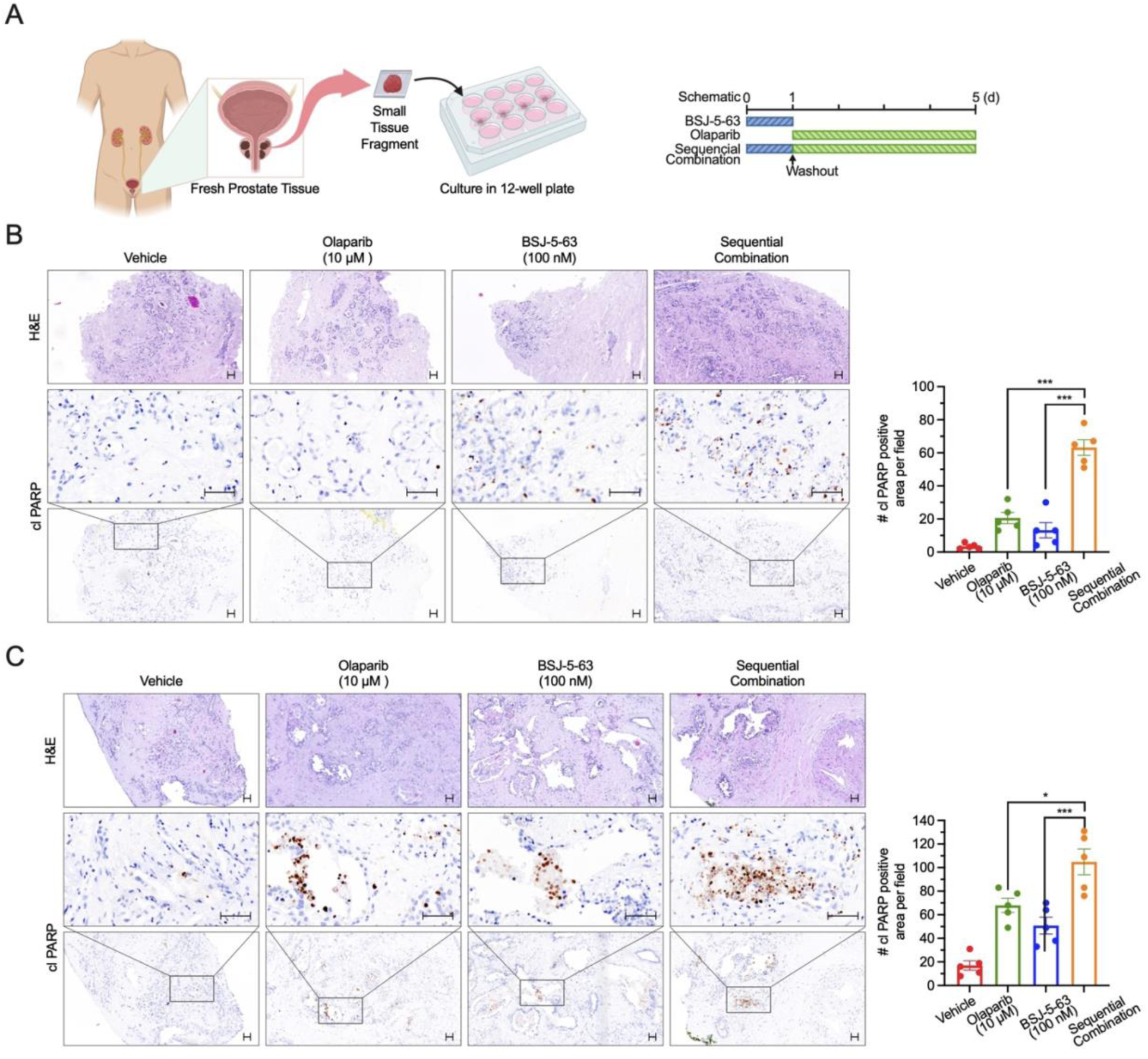
Short-term *ex vivo* antitumor efficacy of sequential combination therapy in PCa tissues from patients #2 and #3. (A) Schematic diagram of the experimental procedure with sequential treatment protocol. (B) Short-term *ex vivo* cultures of PCa tissue fragments from patient #2 (Gleason Score 7a / ISUP grade group 2). Left: Immunohistochemistry analysis and H&E staining showing apoptosis levels in PCa tissues from short-term *ex vivo* cultures. Scale bar: 50 μm. Right: Quantification of cl-PARP staining. Data are presented as mean ± SEM. \*\*\**P* < 0.001; one-way ANOVA with Tukey’s multiple comparisons test. (C) Short-term *ex vivo* cultures of PCa tissue fragments from patient #3 (Gleason score 9 / ISUP grade group 5). Left: Immunohistochemical analysis and H&E staining showing apoptosis levels in PCa tissues from short-term *ex vivo* cultures. Scale bar: 50 μm. Right: Quantification of cl-PARP staining. Data are presented as mean ± SEM. \**P* < 0.05, \*\*\**P* < 0.001; one-way ANOVA with Tukey’s multiple comparisons test.

